# A Dynamic Stem Cell State Links Environmental Cues to Skeletal and Regenerative Fates

**DOI:** 10.64898/2026.11.04.764000

**Authors:** Yuting Wang, Liming Zhao, Kun Chen, Jinyi Xiang, Ethan J. Hunt, Jan Sokol, Ke Ding, Holly M. Steininger, Nathan R. Martinez, Kelly C. Weldon, Edward Chan, David A. Morales, Chao Ma, Sicong Wang, Gabriel Marquez-Arreguin, Honglin Chen, Malachia Y. Hoover, Wan-Jin Lu, Jun Xiao, Fengjing Guo, Michael T. Longaker, Irving L. Weissman, Charles K.F. Chan, Thomas H. Ambrosi

## Abstract

Stem cells faithfully generate tissue architecture during development, yet how their identity and regenerative output are reshaped as anatomical niches mature and environmental signals change with aging and injury remains poorly understood. Rare skeletal stem cells (SSCs) across diverse skeletal sites are defined by combinations of cell-surface markers^1–5^, precluding their genetic interrogation in vivo^6,7^. Here, using spatially informed single cell transcriptomics, we identify Pannexin-3 (Panx3) to mark a dynamic functional state within bona fide SSCs and establish Panx3-Cre^ERT2^ mice for its longitudinal interrogation. Panx3⁺ SSCs exhibit long-term clonal persistence, self-renewal and multipotency, whereas Panx3⁻ SSCs can acquire this state. Successive SSC cohorts engage the Panx3⁺ state during skeletal development, osteoanabolic stimulation expands pre-existing Panx3⁺ SSC clones and injury induces it locally in previously unlabeled progenitors. Thus, anatomically distinct progenitors can converge on a common stem cell state in response to different environmental inputs. Three-dimensional reconstruction of chondrogenic niches further identifies signals that redirect Panx3-lineage cells towards stable cartilage, enabling durable regeneration in aged and osteoarthritic joints and with human SSCs. These findings establish anatomical origin, stem cell state and lineage fate as separable dimensions of stem cell identity and provide genetic access for interrogating and directing endogenous SSC function.

## Introduction

Stem cells are defined by phenotypic identity and functional potential, yet how they transition between functional states in vivo and how these states interact with the local environment to determine lineage output remain fundamental questions in stem cell biology. Development and injury can engage related stem and progenitor populations but produce markedly different tissue outcomes. This suggests that stem cell state and subsequent fate are not necessarily coupled. The emerging diversity of skeletal stem cells (SSCs) provides a rigorous system to address these questions^3,4^. Bona fide mouse and human SSCs have been prospectively isolated using combinations of cell-surface markers and functionally validated by self-renewal and multilineage formation of cartilage, bone and stroma in serial transplantation assays^1,2,6,8,9^. However, reliance on combinatorial surface phenotypes has precluded direct genetic access to these rare cells in their endogenous environment. Consequently, how SSCs are spatially organized, transition between functional states and translate these states into lineage output remains unresolved.

Increasing evidence suggests that skeletal stem and progenitor cell activity is highly heterogeneous across space and time. Pthrp⁺ and FoxA2⁺ populations within the postnatal growth plate exhibit long-term stem cell properties^10,11^, while distinct stem and progenitor cell (SSPC) populations occupy periosteal, endosteal, perivascular, craniofacial and vertebral niches^4,5,12–15^. Although cells labeled by Osterix⁺ and LepR⁺ are widely used to study SSPCs, their established cellular heterogeneity limits their specificity for interrogating bona fide SSCs^16,17^. Human SSCs similarly exhibit distinct lineage potential and clonal behavior across skeletal sites^3^. The identification of multiple stem cell populations within the human growth plate further underscores substantial diversity even within a discrete niche^18^. These findings have largely been interpreted through anatomically or molecularly defined cell populations. This leaves open the possibility that distinct progenitors can converge on shared functional states in response to developmental, systemic and regenerative cues. Addressing this distinction requires genetic access to a functionally validated stem cell population for longitudinal interrogation in vivo, an approach that has remained unavailable for prospectively defined bona fide SSC.

Endochondral ossification, central to skeletal formation and repair, requires SSCs and their progeny to balance stem cell maintenance with sequential chondrogenic, hypertrophic and osteogenic output^8^. In contrast, articular cartilage (AC) maintains a stable non-hypertrophic identity but has limited regenerative capacity. Resident Prg4⁺ and Procr⁺ superficial cells contribute to AC homeostasis and repair^19,20^. However, the persistence of poor intrinsic AC regeneration despite the presence of these resident progenitor-like populations suggests that their activity alone is insufficient to restore substantial cartilage loss, particularly in aging and osteoarthritis (OA). Our previous work established that microfracture (MF) activates phenotypically defined SSCs and that their differentiation can be experimentally directed toward articular cartilage^21^. How injury engages distinct SSC populations and how local environmental signals direct their subsequent lineage output are not fully understood.

Pannexin-3 (Panx3) is expressed in pre-hypertrophic cartilage and has been implicated in chondrocyte and osteoprogenitor differentiation, with genetic loss perturbing endochondral ossification and skeletal growth^22–24^. More recently, Panx3 was shown to be induced during fracture repair and its loss impaired callus formation and bone regeneration^25^. Panx3 is also upregulated in injured and osteoarthritic cartilage but demonstrates context-dependent effects on disease progression^26,27^. These observations suggest that Panx3 may mark a skeletal program engaged across developmental and regenerative settings rather than a fixed differentiated population. Its potential to provide genetic access to bona fide SSCs, distinguish fixed cellular identities from acquirable functional states and reveal how environmental signals shape subsequent lineage fate has not been previously established.

Here, we develop an approach for three-dimensional reconstruction of live skeletal niches linked to single cell transcriptomes and identify Panx3 as marker of a dynamic state within bona fide SSCs shared across developmental and regenerative environments. We further establish Panx3-Cre^ERT2^ mice for genetic interrogation of this state in situ. We demonstrate that distinct skeletal progenitors can engage a common Panx3⁺ stem cell-associated state in response to developmental, systemic and regenerative signals while cells occupying this state retain long-term clonal and multipotent activity. Spatial reconstruction of native skeletal niches further identifies environmental signals that redirect the subsequent lineage output of Panx3-lineage cells. Together, our findings establish anatomical origin, stem cell state and subsequent lineage fate as separable dimensions of stem cell identity, and provide genetic access to bona fide SSCs for investigating skeletal development, aging, disease and regeneration.

## Results

### Spatial single cell mapping identifies a Panx3⁺ stem cell-associated state across distinct niches

To identify cellular states associated with distinct regenerative environments, we compared three chondrogenic skeletal niches with divergent stem and progenitor cell activity and lineage output. Growth plate (GP) and fracture callus (FX) harbor active stem and progenitor populations and undergo endochondral ossification, whereas healthy articular cartilage (AC) has limited SSC activity and regenerative capacity while maintaining a stable non-hypertrophic identity^21,28^. We therefore established a workflow to isolate three-dimensional microexplants from these sites in tamoxifen-induced Actin-Cre^ERT^/Rainbow mice during early postnatal development or following fracture injury (**Fig.1a,b**). Following confocal z-stack imaging, we dissociated freshly isolated microexplants into single cell solution for plate-based, index-sorted whole transcriptome single cell RNAseq analysis (scRNAseq). Unbiased Leiden clustering separated cells into six main subtypes with distinct representations across investigated sites (**Fig.1c-f & Extended Data Fig.1a-c**). All clusters were enriched in genes related to cartilage biology and extracellular matrix formation (**Extended Data Fig.1d,e**). As expected, the GP micro-explant contained chondrogenic resting zone cells (RZ; *Apoe*, *Btg2*) and proliferating/pre-hypertrophic (PZ/PHZ; *Epyc*, *Matn1*) chondrocytes. Hypertrophic chondrocytes were not detected, likely because cartilage microexplants were deliberately collected from the upper GP and because hypertrophic cells may have been preferentially lost during processing due to their larger size. Interestingly, the GP also contained hyaline cartilage-like cell types (*Sdc4*, *Prg4*) mainly found in AC microexplants. Conversely, AC contained cells with characteristics of the GP RZ, suggesting that these anatomically distinct sites share related chondrogenic populations occupying different functional states.. FX microexplants uniquely harbored an endochondral cartilage cell type (*Col10a1*, *Mmp13*) resembling hypertrophic chondrocytes as well as fibrocartilage (*Col3a1*, *Fn1*) known to be important for secondary fracture healing.

**Figure 1:**
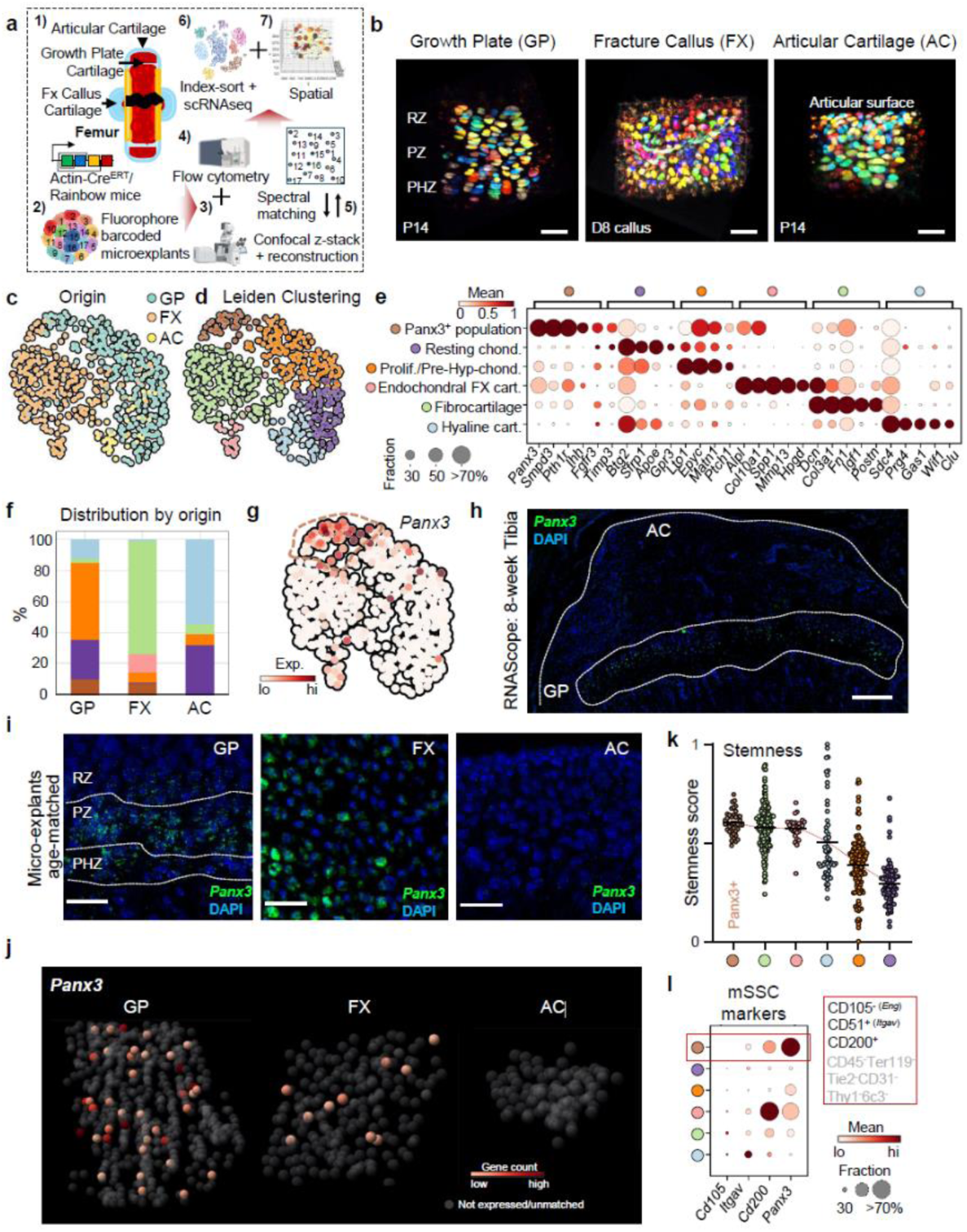
Spatial single cell mapping identifies a Panx3⁺ stem cell-associated state across skeletal niches. **(a)** Schematic overview of the spatial single cell transcriptomic workflow integrating three-dimensional imaging, flow cytometric index-sorting and single cell RNA sequencing. **(b)** Representative confocal z-stacks of microdissected cell microexplants isolated from growth plate (GP), articular cartilage (AC) and fracture callus (FX) of Actin-Cre^ERT^/Rainbow mice for SmartSeq2 scRNAseq. GP, AC tamoxifen induction at P3, harvested 10 days later. FX tamoxifen at 5-month-old and injury 4-weeks later, FX day-8 (D8) analyzed. Scale bars: 30um (GP), 60um (FX), 30um (AC). **(c)** UMAP representation of scRNAseq data colored according to the microexplants’ origin. **(d)** Unbiased Leiden clustering resolving six major chondrogenic populations. **(e)** Dot plot of representative marker genes defining the six Leiden clusters. **(f)** Cross-tabulation of Leiden cluster identity by tissue of origin. **(g)** UMAP feature plot of Panx3 expression, identifying preferential enrichment within the Panx3+ cluster population. **(h)** RNAscope analysis of tibiae from 8-week-old mice validating the spatial restriction of Panx3 expression to the GP. Scale bars: 300um. **(e)** RNAscope analysis of age-matched cartilage tissues corresponding to the sequenced micro-explants, confirming the tissue-specific distribution of Panx3 expression. Scale bars: 3um. **(j)** Three-dimensional reconstruction of representative microexplants illustrating the spatial distribution of Panx3 expression across distinct cartilage tissue origins. **(k)** Inferred stemness scores of individual cells stratified by Leiden clusters. **(l)** Dot plot of Panx3 and canonical SSC markers across Leiden clusters.

Across these transcriptionally related chondrogenic populations, Pannexin-3 (Panx3) distinguished a discrete cellular state present in the stem and progenitor cell-active GP and FX niches but absent from AC. Panx3^+^ cells co-expressed a combination of classic chondrocyte markers (*Sox9*, *Acan*, *Col2a1*) with varying levels of RZ and PZ genes as well as particular enrichment for PHZ markers such as *Epyc, Pth1r* and *Ihh* (**Fig.1e,g & Extended Data Fig.1d**). In line, gene enrichment analysis for GO Biological Processes suggested an important role for Panx3 cluster cells in skeletal system development, response to hypoxia and negative regulations of apoptosis (**Extended Data Fig.1e**). Fittingly, previous reports showed *Panx3* preferentially labeling cells of the PHZ in cartilage anlagen during development and in the GP^22,24,29^. RNAScope-based in situ hybridization similarly and selectively detected *Panx3* RNA in the 8-week-old murine tibial growth plate but not at AC, periosteal, endosteal or bone marrow sites (**Fig.1h & Extended Data Fig.1f**). To confirm the location of Panx3-expressing cells more precisely, we performed RNA in situ hybridization at microexplant-matched sites (**Fig.1i**). Here, we found a spatial gradient of Panx3 mRNA across the GP, with progressively increasing expression from the lower RZ toward the PHZ zone. Confirming scRNAseq data, Panx3 expression was absent from AC.

To further link single cell transcriptomes with three-dimensional spatial information, we developed a workflow leveraging the Rainbow reporter to barcode individual cells within intact microexplants (**Fig. 1a**; see Methods). Confocal imaging captured the spatial position and four-channel fluorescence profile of each cell before dissociation and FACS index-sorting for Smart-seq2^30^. An image analysis pipeline segmented cells in confocal z-stacks and quantified cell size and fluorescence intensity. Matching normalized fluorescence profiles between imaged cells and FACS events enabled assignment of spatial coordinates to individual transcriptomes and single cell spatial reconstruction. The results spatially resolved chondrogenic gene expression patterns across the three cartilage sites (**Extended Data Fig.1g**) and established a framework for subsequent interrogation of cellular crosstalk within endogenous three-dimensional niches. Notably, the approach validated distinct *Panx3* expression across RZ to PHZ and in FX callus, but not in AC, reinforcing the association of Panx3 with niches exhibiting active skeletal stem/progenitor activity (**Fig.1j**).

Together, the restricted localization of Panx3⁺ cells to stem cell-active environments, their transcriptional heterogeneity and their positioning along the GP axis prompted us to ask whether Panx3 marks a skeletal stem/progenitor population. Supporting this possibility, comparison with our published transcriptomic data of prospectively isolated SSC lineage populations revealed *Panx3* expression restricted to phenotypic SSCs as well as their immediate multipotent bone-cartilage-stromal progenitors (BCSPs) and no expression in more committed populations (**Extended Data Fig.1h**)^14,31^. We therefore examined the inferred stemness of the six chondrogenic populations using validated algorithms^32^. To that end our approach not only considered developmental potential from transcriptional diversity but also the log-transformed number of genes detected per cell as a direct measure of transcriptional complexity. Indeed, the Panx3 cluster showed the highest stemness score (**Fig.1k**). This was further matched by the enrichment of canonical SSC markers within our Panx3 cluster (**Fig.1l**). Notably, the RZ supopulation showed comparatively the lowest stemness score and SSC marker expression despite the established presence of stem/progenitor activity within these regions. Consistent with recent findings identifying transcriptionally distinct stem-like populations within the RZ of the human GP^18^, these data potentially highlight heterogeneity within the GP and suggest that stem-like activity resides within discrete subpopulations rather than uniformly distributed across canonical chondrocyte states. Notably, *Panx3* cells did not overlap with the recently identified Procr⁺ chondrogenitors of the AC superficial zone^20^, which emerge around postnatal week 4 in mice (**Extended Data Fig.1d & 2a,b**). Consistent with our mouse findings, *PANX3*⁺ cells were preferentially detected in human embryonic endochondral bone relative to cranial bone and were similarly concentrated across the RZ-to-PHZ in adolescent GPs (**Extended Data Fig.2c-f**)^33^. Altogether, these findings identify Panx3 as a candidate marker of a stem cell-associated state selectively engaged within chondrogenic niches with active SSC function.

### Panx3 provides genetic access to prospectively defined phenotypic SSCs

Mouse SSCs have previously required prospective isolation using combinations of cell-surface markers, limiting their direct genetic interrogation in vivo^7^. To determine whether Panx3 could provide genetic access to this functionally defined stem cell population, we generated tamoxifen-inducible Panx3-Cre^ERT2^ mice and crossed them with either Rainbow or Ai9/tdTomato reporter mice to define the in vivo dynamics and lineage behavior of Panx3⁺ cells (**Fig.2a & Extended Data Fig.2g**; see Methods). We observed no reporter leakage in the absence of tamoxifen (**Extended Data Fig.2h**). The Rainbow reporter enables simultaneous visualization of clonally related cells through stochastic, heritable expression of distinct fluorescent proteins, allowing assessment of clonal expansion and persistence in situ. Given that clonal expansion is a stem cell hallmark, we first analyzed GPs of Panx3-Cre^ERT2^/Rainbow mice. A one-week pulse-chase initiated at postnatal day 3 (P3) revealed distinct clones throughout the GP and developing secondary ossification center (**Fig.2b**). Clones ranged from individual labeled cells to expanded columnar/multi-cellular populations, consistent with different cellular activity. Notably, the capacity for clonal expansion of Panx3+ cells persisted into adulthood and remained detectable in GP remnants of 16-month-old mice (**Fig.2c,d**). To further define the proliferative behavior of Panx3⁺ GP cells, we combined the Ai9/tdTomato reporter with EdU labeling at neonatal and adult stages. In P12 mice induced with tamoxifen at P3, a short 6h EdU pulse revealed preferential EdU incorporation in Panx3-lineage cells within the PZ, whereas a subset of tdTomato⁺ cells located closer to the RZ remained EdU^-^(**Fig.2e**), indicating heterogeneous proliferative activity within neonatally labeled Panx3-lineage cells. In a separate experiment in 3-month-old mice, concurrent tamoxifen and 5-day EdU labeling revealed EdU⁻ tdTomato^+^ cells positioned above EdU⁺ tdTomato⁺ columnar populations, supporting the coexistence of relatively quiescent and proliferative Panx3⁺ cell states within the adult GP. This pattern was consistent with the distinct activity states of individual Panx3^+^ clones observed by Rainbow reporter analysis. Together, these findings demonstrate heterogeneous proliferative activity among Panx3-lineage cells and long-term persistence of the Panx3-marked population.

**Figure 2:**
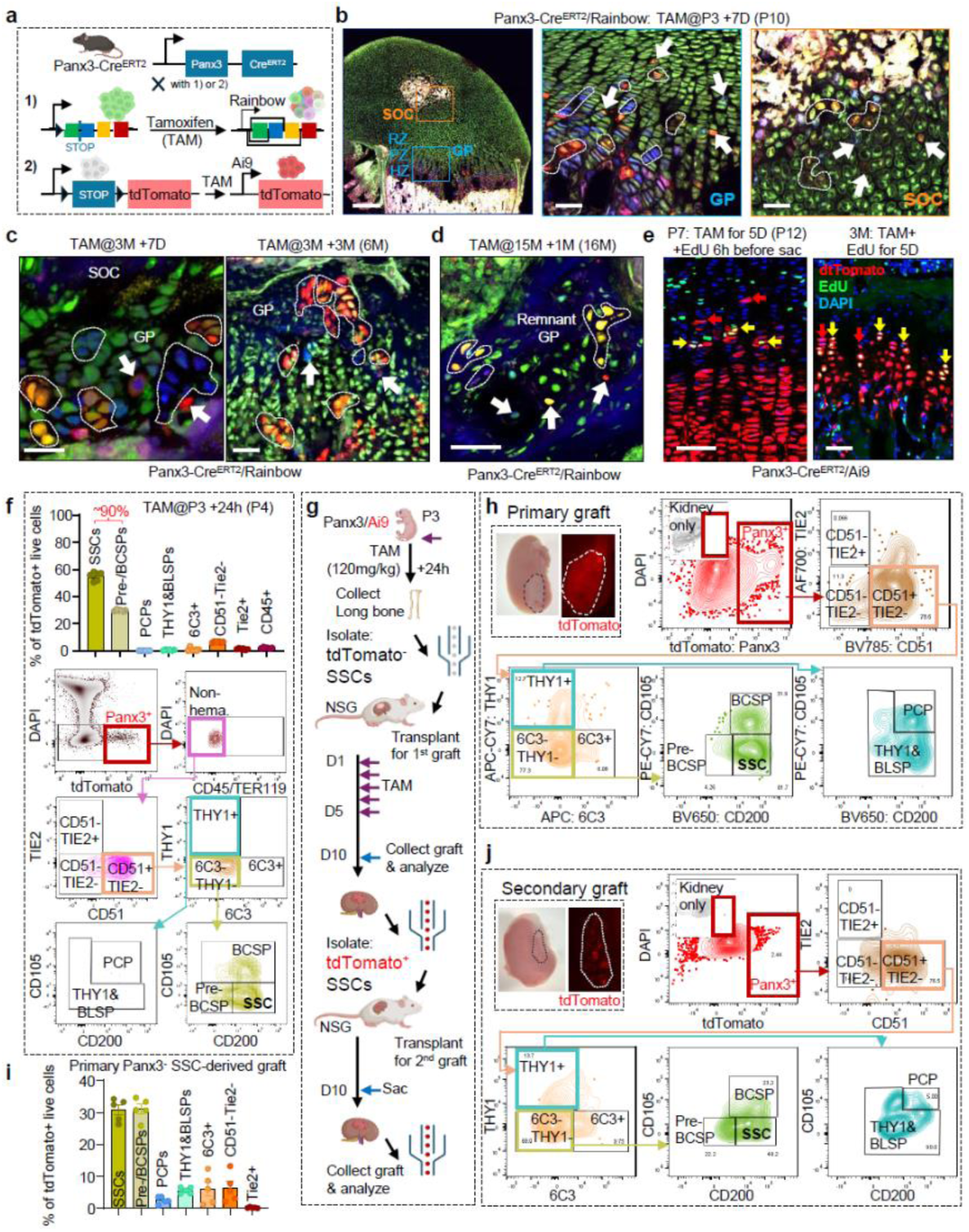
Panx3 labels bona fide skeletal stem cells. **(a)** Schematic of the breeding strategy used to generate Panx3-Cre^ERT2^/Rainbow and Panx3-Cre^ERT2^/Ai9 mice for lineage tracing and reporter analysis. **(b)** Clonal labeling and expansion within the secondary ossification center (SOC) and GP of Panx3-Cre^ERT2^/Rainbow mice induced with tamoxifen at P3 and analyzed 7 days later. Scale bars: 300um (left), 30um (middle, right). **(c)** Clonal labeling and expansion of Panx3-lineage cells within the growth plate of adult Panx3-Cre^ERT2^/Rainbow mice induced with tamoxifen at 3 months of age and analyzed after 7 days (left) or 3 months (right). Scale bars: 30um. **(d)** Persistent clonal labeling and expansion of Panx3-lineage cells within growth plate remnants of aged Panx3-Cre^ERT2^/Rainbow mice induced with tamoxifen at 15 months of age and analyzed 1 month later. Scale bars: 50um. **(e)** EdU labeling of Panx3-lineage cells in the distal femoral GP of Panx3-Cre^ERT2^/Ai9 mice. Left, mice were induced with tamoxifen at P3 and analyzed at P12, with EdU administered 6h before charvest. Right, 3-month-old mice received tamoxifen induction and EdU labeling for 5 consecutive days before harvest. Scale bars: 30um. **(f)** Quantification of phenotypically defined SSC populations among tdTomato⁺ live cells (top) and representative flow cytometry plots (bottom) from P3 mice analyzed 24h after tamoxifen induction (0.3 mg/pup). *n* = 6 biologically independent samples. **(g)** Schematic of the serial renal capsule transplantation assay used to assess the stem cell properties and lineage plasticity of Panx3⁺ SSCs isolated from P4 mice. **(h)** Brightfield and fluorescence images of primary renal capsule grafts following tamoxifen induction, together with representative flow cytometric analysis of graft-derived tdTomato⁺ cells. *n* = 5 biologically independent samples. **(i)** Quantification of phenotypically defined SSC populations among tdTomato⁺ live cells recovered from primary renal capsule grafts. *n* = 5 biologically independent samples. **(j)** Brightfield and fluorescence images of secondary renal capsule grafts, together with representative flow cytometric analysis of graft-derived tdTomato⁺ cells.

We previously defined mouse SSCs and their downstream lineage populations through prospective isolation using combinatorial cell-surface markers (**Extended Data Fig.1i**)^1^. We therefore directly tested whether Panx3-Cre^ERT2^ labeling genetically accesses this prospectively defined SSC hierarchy. Following a 24h tamoxifen pulse, flow cytometric analysis of long bones revealed striking selectivity of Panx3 labeling for phenotypic SSCs (CD45^-^Ter119^-^Tie2^-^Thy1^-^6C3^-^CD51^+^CD200^+^), with the second highest labeling observed in multipotent pre-BCSP/BCSP populations together accounting for ∼90% of labeled cells and minimal labeling among more committed descendants (**Fig.2f**). These results indicated that Panx3-Cre^ERT2^ provides highly selective genetic access to the prospectively defined SSC compartment and its earliest progeny, bridging the established surface marker definition of bona fide SSCs with longitudinal interrogation in vivo. Conversely, a substantial fraction of phenotypic SSCs lacked Panx3-tdTomato lineage labeling, which likely reflects incomplete recombination and phenotypic SSCs in non-GP regions (i.e., periosteum), but could also explain distinct SSC activity states (**Extended Data Fig.2i,j**). Panx3 restriction to the lower RZ further distinguished Panx3⁺ SSCs from other GP stem cell populations. Pthrp⁺ RZ cells partially overlap with phenotypically defined SSCs and exhibit long-term stem cell activity^10^, supporting functional diversity within this compartment. These findings suggested that Panx3 marks a dynamic state within the phenotypically defined SSC compartment. Hereafter, we use Panx3⁺ SSCs to refer specifically to prospectively defined SSCs expressing or labeled by Panx3, whereas the Panx3⁺ state denotes the Panx3-associated cellular program that can be acquired by skeletal progenitors.

We next asked whether Panx3 identifies a stable SSC subtype or an acquirable state within the SSC compartment. Following a one-time, high-dose (120mg/kg) tamoxifen-labeling for 24h in Panx3-Cre^ERT2^/Ai9 mice to maximize recombination, we prospectively isolated tdTomato^-^phenotypic SSCs from long bone GPs and transplanted them beneath the renal capsules of NSG recipient mice (**Fig.2g**). We reasoned that transplantation into an ossicle formation-inducing niche might activate previously Panx3⁻ SSCs and induce Panx3 expression. Indeed, with tamoxifen administered during the first five days after transplantation, grafts generated from initially tdTomato⁻ SSCs contained tdTomato⁺ SSCs together with the full spectrum of downstream SSC populations. Thus, prospectively defined Panx3⁻ SSCs can acquire the Panx3⁺ state while retaining the capacity to reconstitute the broader SSC lineage (**Fig.2h,i**). To assess long-term stem cell activity, we prospectively isolated tdTomato^+^/Panx3⁺ SSCs from primary grafts and performed secondary transplantation. These cells reproducibly generated grafts containing all major SSC lineage populations, demonstrating self-renewal and multilineage differentiation capacity (**Fig.2j**). In total, these experiments establish Panx3-Cre^ERT2^ as a genetic reporter of bona fide SSCs while demonstrating that Panx3 does not define a fixed SSC subtype. Instead, it marks a dynamic state that can be acquired within the prospectively defined SSC compartment while retaining clonal self-renewal and multilineage potential.

### Successive waves of Panx3⁺ stem cell activity support skeletal development and maintenance

The rarity of SSCs and lack of tools to trace these cells in their endogenous environment have limited our understanding of their spatial distribution and lineage contributions in situ. To qualitatively and quantitatively define Panx3⁺ cell behavior, we first analyzed Panx3-Cre^ERT2^/Ai9 mice during embryonic development. TdTomato⁺ cells were detected between E10.5 and E12, labeling much of the developing primary ossification center (**Fig.3a,b & Extended Data Fig.3a**). Lineage tracing initiated at E10.5 revealed contributions to cartilage, bone and stromal compartments, marked by Aggrecan, Collagen-I and Cxcl12, respectively (**Fig.3c & Extended Data Fig.3b**). Notably, embryonically labeled Panx3⁺ cells progressively diminished and were largely absent by early postnatal life (**Fig.3b,d**). A second tamoxifen pulse at birth marked a distinct wave of tdTomato⁺ cells and sequential labeling at E10.5 and P0 marked nearly the entire primary ossification center by P3 (**Fig.3d & S3c**). This pattern resembles the previously described contribution of Osx-lineage cells with the distinction that Panx3 lineage cells may occupy a higher position in the cellular hierarchy, providing the endochondral template from which Osx⁺ bone forming cells emerge^16^.

**Figure 3:**
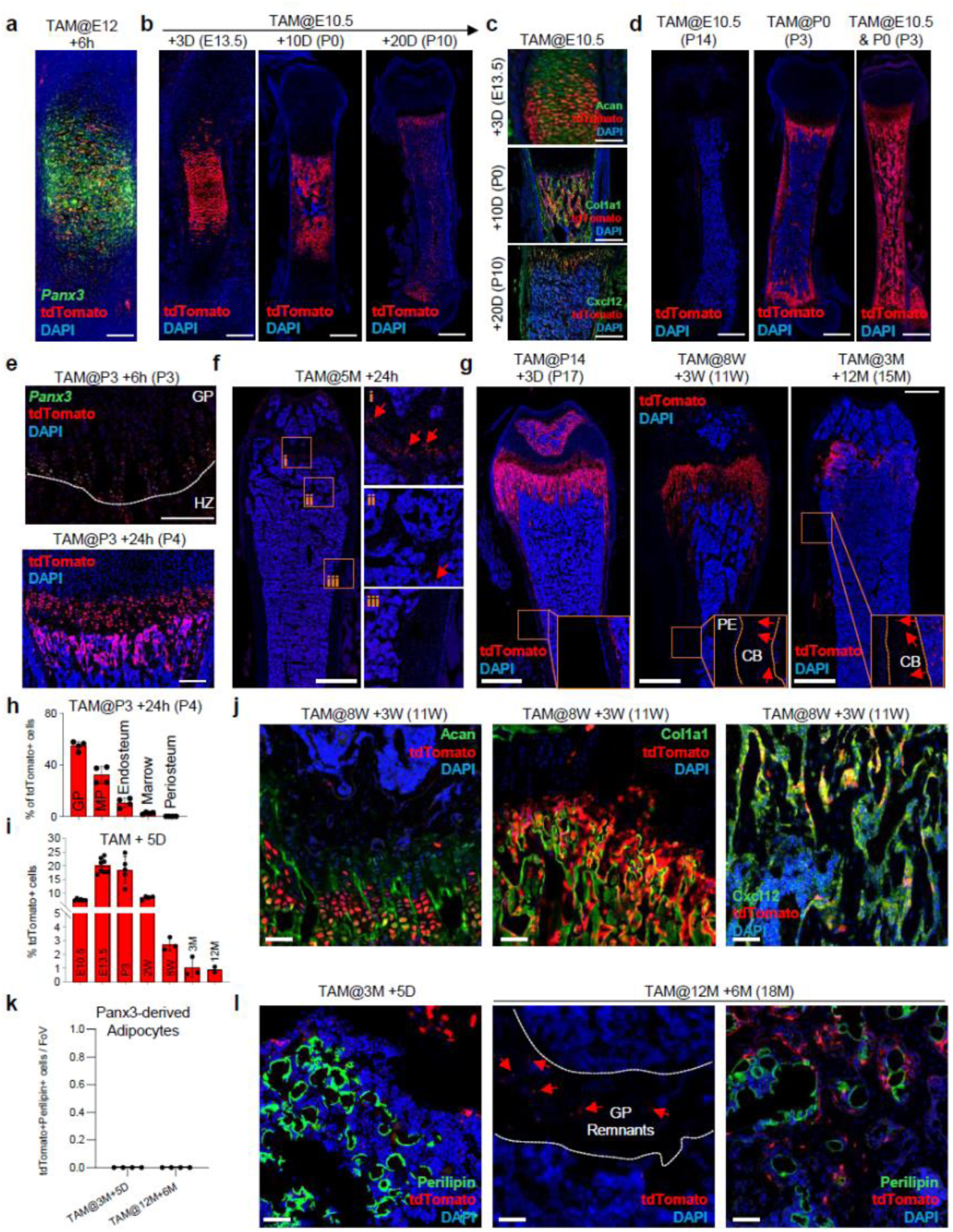
Successive waves of Panx3⁺ SSC activity shape skeletogenesis. **(a)** Representative E12 embryonic femur of Panx3-Cre^ERT2^/Ai9 mouse tamoxifen-induced and analyzed 6h later. Co-stained using an RNAscope probe against *Panx3*. Scale bars: 20um. **(b)** Representative femur sections of Panx3-Cre^ERT2^/Ai9 induced at E10.5 and followed for indicated timeframe. Scale bars:40um (e13.5), 0.4mm (P0), 0.5mm (P10). **(c)** Representative co-staining of E10.5 induced Panx3-Cre^ERT2^/Ai9 mouse femurs co-stained for Aggrecan (Acan), Collagen-1a1 (Col1a1) and Cxcl12. Scale bars: 0.2um (e13.5), 0.4mm (P0, P10). **(d)** Representative femur sections of Panx3-Cre^ERT2^/Ai9 induced at indicated indicated timepoints. Scale bars: 0.5mm (p14), 0.4mm (P0, P10). **(e)** Representative P3 femur of Panx3-Cre^ERT2^/Ai9 mouse tamoxifen-induced and analyzed 6h (top) or 24h (bottom) later. Top: Co-stained using an RNAscope against Panx3. Scale bars: 80um. **(f)** Representative femur sections of Panx3-Cre^ERT2^/Ai9 induced at 5-months of age traced for 24h. GP, metaphyseal and periosteal regions size-enhanced on the right. Scale bars: 2mm. **(g)** Representative femur sections of Panx3-Cre^ERT2^/Ai9 induced at different indicated timepoints and traced for short- and long-term periods. Scale bars: 2mm. **(h)** Quantification of Panx3-derived tdTomato+ cells in distinct femoral subregions (MP, metaphysis) after 24 chase at P3. *n* = 4 biologically independent samples. **(i)** Quantification of Panx3-derived tdTomato+ cells at different timepoints across life. *n* = 2-9 biologically independent samples. **(j)** Representative co-staining of 8-week-old induced Panx3-Cre^ERT2^/Ai9 mouse femurs co-stained for Acan, Col1a1 and Cxcl12. Scale bars: 50um. **(k)** Quantification of Panx3-derived Perilipin+ bone marrow adipocytes. *n* = 4 biologically independent samples. **(l)** Corresponding representative co-staining of Panx3-Cre^ERT2^/Ai9 mouse femurs co-stained for Perilipin. Scale bars: 50um.

We next examined the kinetics of the Panx3-lineage postnatally. A 6h chase in P3 mice labeled a small population restricted to the lower RZ, PZ and PHZ, recapitulating the spatial distribution observed by *Panx3* RNA detection (**Fig.1 & 3e**). By 24h, labeled cells had expanded into columns and extended into the primary ossification center, raising the possibility that a subset of GP progenitors transitioning toward hypertrophy subsequently contributes to the osteogenic compartment. In skeletally mature 5-month-old mice, a 24h pulse-chase labeled columnar GP cells as well as individual cells along trabecular surfaces, reflecting the reduced activity of the mature GP and suggesting establishment and persistence of a potentially latent Panx3^+^ skeletal progenitor reservoir outside the GP (**Fig.3f**). Longer term tracing initiated at P14, 8-weeks, 3-months or 12-months further demonstrated ongoing Panx3-lineage contribution to metaphyseal, endosteal and cortical bone, but not the periosteum (**Fig.3g & Extended Data Fig.3d**). The GP remained the area most enriched in Panx3^+^ cells in postnatal life (**Fig.3h & Extended Data Fig.3e**). Combined, the abundance of tdTomato⁺ cells was highest early in life and progressively declined with age, paralleling the reduction in SSC activity upon maturation (**Fig.3i**). Nevertheless, Panx3-lineage cells continued to contribute to mature skeletal tissues throughout adulthood (**Fig.3j & Extended Data Fig.3f**). The decline of embryonically labeled cells together with the emergence of a distinct neonatal Panx3⁺ wave revealed successive periods of Panx3⁺ SSC engagement during skeletogenesis, while subsequent lineage tracing demonstrated that individual Panx3⁺ SSCs can persist long term and sustain clonal, multilineage skeletal output.

Skeletal aging has traditionally been viewed through the lens of an osteogenic versus adipogenic bifurcation choice from a shared mesenchymal progenitor. The persistence of Panx3-lineage cells into adulthood allowed us to test this paradigm in a genetically traced SSC population. Consistent with our previous findings that mouse and human SSCs do not generate marrow adipocytes^1,2,14^, Panx3-lineage cells remained absent from Perilipin⁺ adipocytes with aging (**Fig.3k,l**). These findings further challenge the traditional osteogenic versus adipogenic lineage shift model of skeletal aging, arguing against an age-dependent fate bias within a common SSC population.

Finally, Panx3-driven tdTomato labeling also extended to vertebral growth plates (**Extended Data Fig.4a**). Surprisingly, Panx3⁺ cells were present in developing mandibles and cranial sutures but diminished following skeletal maturity (**Extended Data Fig.4b,c**). Unlike the endochondral GP, these craniofacial sites were derived from distinct embryonic origins and predominantly undergo intramembranous ossification. Nevertheless, cranial sutures can harbor SSC populations supporting both intramembranous and endochondral bone formation^4^. Panx3⁺ cells therefore extend across distinct developmental and ossification contexts. In line with this, adult mandibular SSCs can reactivate developmental neural crest programs during distraction osteogenesis^34^. Collectively, these lineage-tracing studies reveal successive, anatomically distinct waves of Panx3⁺ cells that generate cartilage, bone and stroma, persist into adulthood without adipogenic contribution and extend across distinct skeletal sites. Their changing distribution across development and maturation further supports Panx3 as a dynamically engaged SSC state rather than a fixed anatomical population.

### The Panx3⁺ state responds dynamically to systemic anabolic stimulation and injury

Because Panx3⁺ cells persisted within the GP and along trabecular surfaces of the adult skeleton (**Fig.3e-h & Extended Data Fig.4d**), we next asked whether the Panx3⁺ state responds to distinct physiological and regenerative stimuli. Intermittent parathyroid hormone (PTH1–34) is a potent osteoanabolic stimulus that has been linked, at least in part, to activation of skeletal progenitor populations^35^. We first assessed clonal behavior in 7-month-old Panx3-Cre^ERT2^/Rainbow mice induced with tamoxifen one week before initiating two weeks of PTH1–34 treatment. PTH1-34 significantly increased the number of Panx3-labeled clones and the number of cells per clone within the GP, indicating activation of local Panx3⁺ SSCs (**Fig.4a**). Similarly, when we treated Panx3-Cre^ERT2^/Ai9 mice with PTH1-34, we observed enhanced Panx3-SSC lineage activity in microdissected epiphyseal/GP and metaphyseal/trabecular femoral regions at the 2-week timepoint (**Extended Data Fig.4e**). Four weeks of PTH1-34 resulted in the expected increase in trabecular bone parameters (**Fig.4b**). The treatment likewise increased the abundance of tdTomato⁺ lineage cells throughout the femur, including the GP, trabecular and cortical compartments (**Fig.4c-e**). Thus, a clinically relevant osteoanabolic stimulus acts on the genetically defined Panx3⁺ SSC compartment, expanding pre-existing clonal activity and subsequent skeletal contribution.

**Figure 4:**
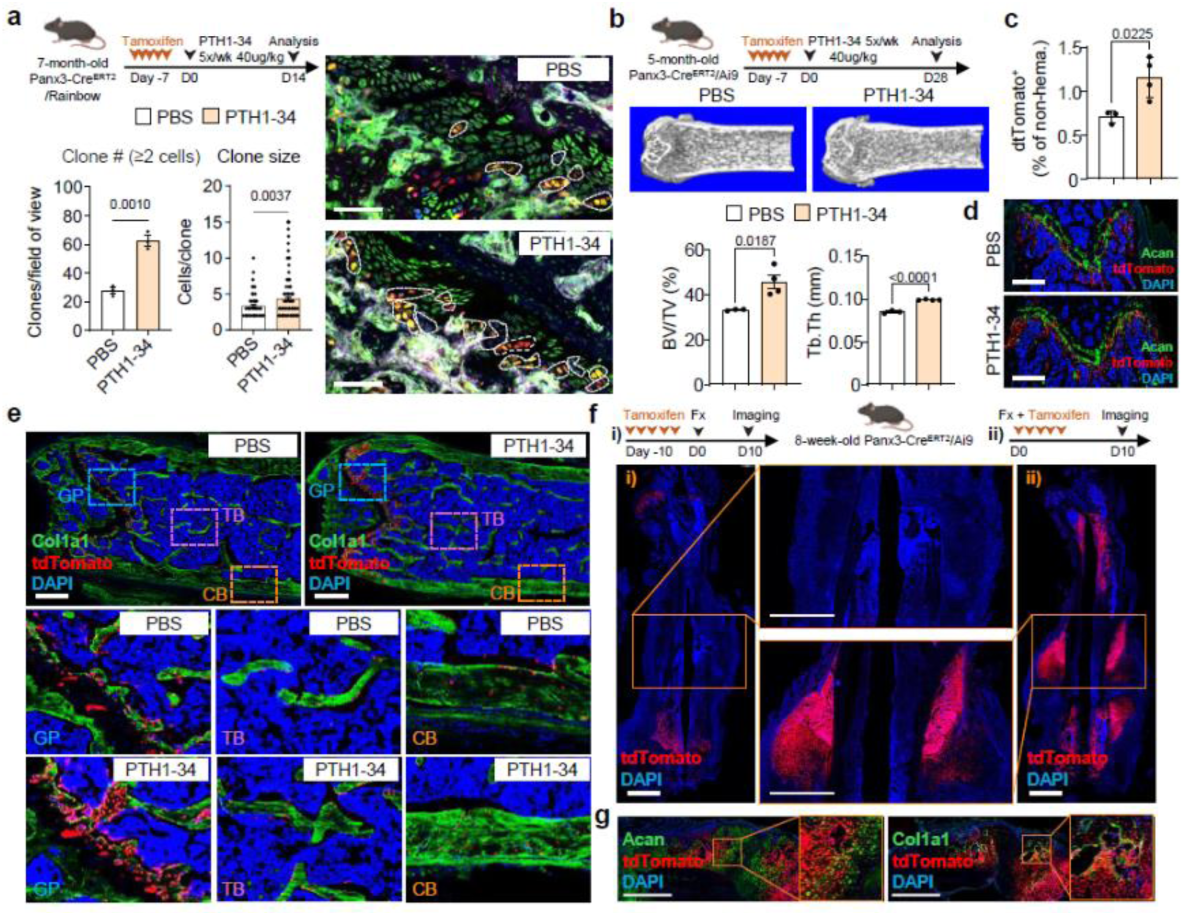
Systemic and local cues differentially engage the Panx3⁺ stem cell state. **(a)** Experimental schematic of 2-week Parathyroid Hormone (PTH1-34) treatment in Panx3-Cre^ERT2^/Rainbow mice for clonal analysis (top). Quantification of total clone number and number of cells per clone (left) as well as representative images thereof (right). Data are presented as mean ± SEM; *n* = 3 biologically independent samples. Statistical significance was assessed using a two-tailed unpaired Student’s *t* test. Exact *P* values are indicated in the figure, Scale bars: 300um. **(b)** Experimental schematic of 4-week Parathyroid Hormone (PTH1-34) treatment in Panx3-Cre^ERT2^/Ai9 mice (top). Representative micro-CT images (middle) and quantification of volumetric bone volume (BV/total volume -TV) as well as trabecular thickness (Tb.Th; right). Data are presented as mean ± SEM; *n* = 3-4 biologically independent samples. Statistical significance was assessed using a two-tailed unpaired Student’s *t* test. Exact *P* values are indicated in the figure. **(c)** Flow cytometric assessment of tdTomato^+^ cells in femurs of control treated or PTH1-34 treated Panx3-Cre^ERT2^/Ai9 mice. Data are presented as mean ± SEM; *n* = 3-4 biologically independent samples. Statistical significance was assessed using a two-tailed unpaired Student’s *t* test. Exact *P* values are indicated in the figure. **(d)** Representative images of tdTomato signal in metaphyseal GP region co-stained with Acan. Scale bars: 1mm. **(e)** Representative images of tdTomato signal in indicated femoral subregions co-stained with Col1a1. Scale bars: 1mm. **(f)** Experimental schematic of bi-cortical fracture experiments in Panx3-Cre^ERT2^/Ai9 mice (top). Representative images enlarged callus region (middle). Scale bars: 2mm. **(g)** Bi-cortical fracture callus region of Panx3-Cre^ERT2^/Ai9 mice tamoxifen-induced at time of injury co-stained with Acan and Col1a1. Scale bars: 2mm.

We next sought to answer whether Panx3⁺ cells contribute to repair following injury. Given their localization to the GP and metaphysis, but not the periosteum (a major source of cells during diaphyseal bi-cortical fracture repair^36,37^), we hypothesized that these locally restricted Panx3⁺ cells may not directly participate in response to cortical injury of the diaphysis. Indeed, labeling initiated 10 days before fracture resulted in no detectable Panx3-lineage contribution to the day-10 fracture callus (**Fig.4f**). In contrast, tamoxifen administration initiated at the time of fracture produced extensive lineage labeling throughout the callus, including both chondrogenic and osteogenic populations (**Fig.4f & Extended Data Fig.4f**). Rather than excluding Panx3⁺ cell involvement in fracture repair, these findings align with the injury-induced plasticity of periosteal SSCs, which shift from intramembranous to endochondral activity after injury^12^. The emergence of Panx3-lineage cells therefore indicates local acquisition of the Panx3⁺ state during injury-induced progenitor activation. Together, these findings reveal distinct modes through which environmental perturbations engage the Panx3⁺ state. Systemic anabolic stimulation expands pre-existing Panx3⁺ SSC activity, whereas injury induces local acquisition of the state in previously unlabeled skeletal progenitors.

### Panx3⁺ cell depletion compromises skeletal development, maintenance and regeneration

To determine the functional requirement for cells occupying the Panx3⁺ state, we generated Panx3-Cre^ERT2^/DTA mice, enabling tamoxifen-dependent ablation of Panx3⁺ cells (**Fig.5a**). Global Panx3-knockout has been reported to cause dwarfism and reduced bone mass, accompanied by expansion of the PZ/PHZ and shortening of the HZ in the GP^24,38^. In line with these phenotypes, embryonic or early postnatal ablation of Panx3⁺ cells at P1 impaired normal skeletal growth and development (**Fig.5b,c & Extended Data Fig.5a,b**). As predicted by our lineage tracing studies (**Fig.2,3**), DTA-mediated ablation resulted in reduced abundance of phenotypic SSCs, conforming with selective targeting of SSCs by Panx3⁺ cell ablation (**Fig.5d & Extended Data Fig.5c**). The modest but significant reduction may reflect replenishment from Panx3⁻ SSCs following tamoxifen clearance. Of note, osteochondrogenic progenitors remained unchanged at this timepoint, whereas 6C3^+^ stromal progenitors increased. This differential response may reflect compensatory expansion of distinct marrow stromal populations, consistent with the presence of multiple stromal progenitor lineages within the bone marrow^6,39^.

**Figure 5:**
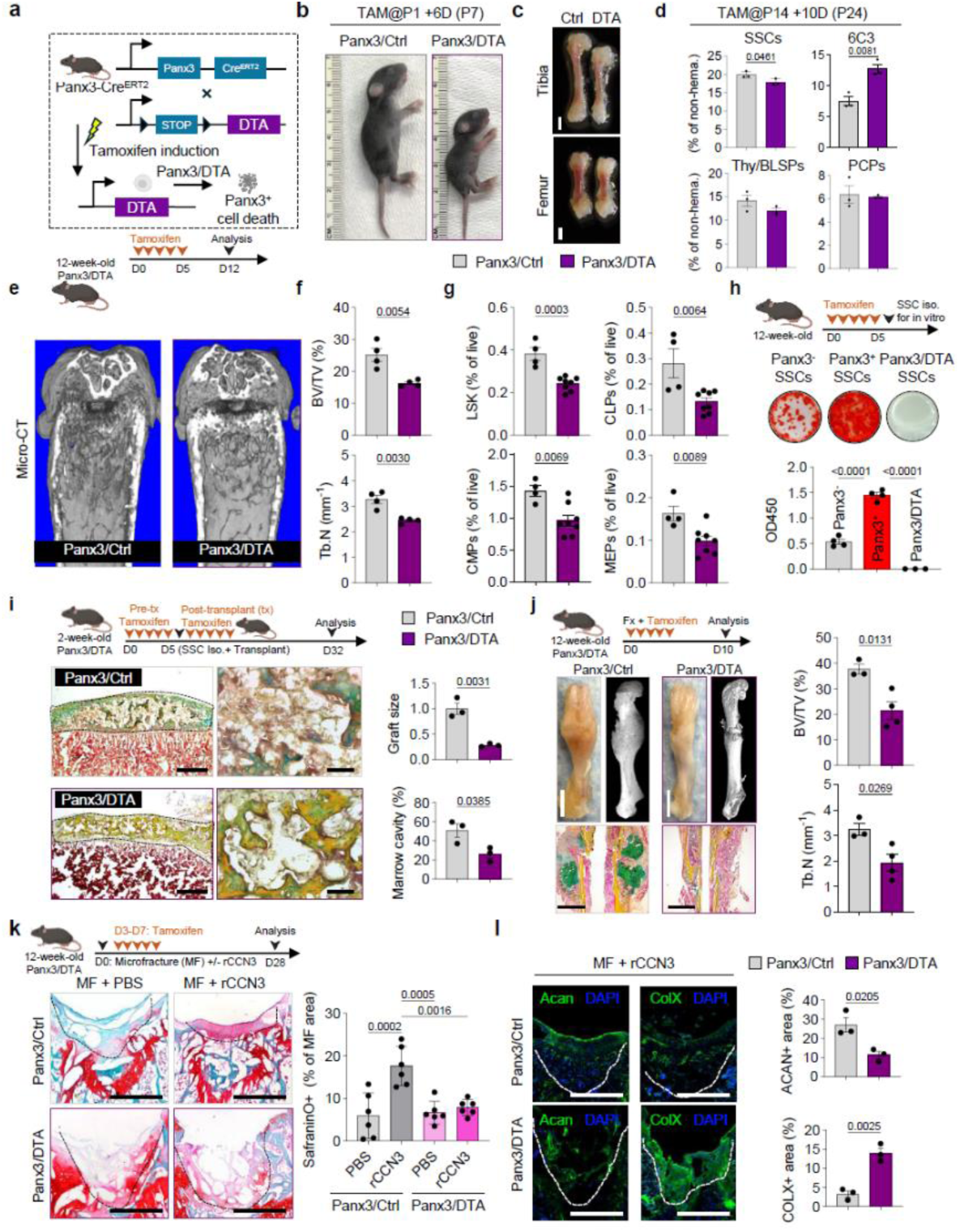
Panx3⁺ SSCs are required for skeletal development, maintenance and regeneration. **(a)** Schematic of the breeding strategy used to generate Panx3-Cre^ERT2^/Diphteria Toxin (DTA) mice to ablate cells expression *Panx3*. **(b)** Representative images of control (Ctrl) or Panx3-Cre^ERT2^/DTA mice induced at P1 and sacrificed at P7. **(c)** Images of corresponding tibia and femur bones. Scale bars: 1mm. **(d)** Flow cytometric analysis of SSC lineage populations of Panx3-Cre^ERT2^/DTA mice tamoxifen-induced at P14 and sacrificed ten days later. Data are presented as mean ± SEM; *n* = 3 biologically independent samples. Statistical significance was assessed using a two-tailed unpaired Student’s *t* test. Exact *P* values are indicated in the figure. **(e)** Experimental schematic of Panx3-Cre^ERT2^/DTA adult mouse analysis (top). Representative micro-CT images of femurs (bottom). **(f)** Quantification of micro-CT data. Data are presented as mean ± SEM; *n* = 4 biologically independent samples. Statistical significance was assessed using a two-tailed unpaired Student’s *t* test. Exact *P* values are indicated in the figure. **(g)** Analysis of bone marrow hematopoietic stem cell lineage populations in same mice. Data are presented as mean ± SEM; *n* = 4-8 biologically independent samples. Statistical significance was assessed using a two-tailed unpaired Student’s *t* test. Exact *P* values are indicated in the figure. **(h)** Prospectively isolated SSCs compared for their in vitro mineralization capacity in presence of an osteogenic cocktail. Data are presented as mean ± SEM; *n* = 4 biologically independent samples. Statistical significance was assessed using a two-tailed unpaired Student’s *t* test. Exact *P* values are indicated in the figure. **(i)** Renal capsule transplantation of prospectively isolated cells from Panx3-Cre^ERT2^/DTA mice. Experimental schematic (top left), representative Movat’s Pentachrome staining of derived grafts (bottom left) and quantification of graft and marro cavity size formed (right). Scale bars: 0.3mm(left), 80um(right). Data are presented as mean ± SEM; *n* = 3 biologically independent samples. Statistical significance was assessed using a two-tailed unpaired Student’s *t* test. Exact *P* values are indicated in the figure. **(j)** Bi-cortical fracture experiment in Panx3-Cre^ERT2^/DTA mice with experimental schematic (top left), representative images of callus formed as photograph, micro-CT image and Movat’s Pentachrome staining (bottom left) and micro-CT based quantification of callus parameters (right). Scale bars:6mm(femur), 2mm(callus). Data are presented as mean ± SEM; *n* = 3-4 biologically independent samples. Statistical significance was assessed using a two-tailed unpaired Student’s *t* test. Exact *P* values are indicated in the figure. **(k)** Microfracture (MF) experiment in Ctrl and DTA mice with or without recombinant CCN3 (rCCN3). Experimental schematic (top left), representative Safranin O staining (bottom left) and quantification of Safranin O^+^ regenerate area (right). Scale bars: 0.4mm. Data are presented as mean ± SEM; *n* = 6 biologically independent samples. Statistical significance was assessed using one-way ANOVA with Tuckey’s multiple comparison test. Exact *P* values are indicated in the figure. **(l)** Representative immunostaining with Acan and Collagen-X (ColX) of MF treated with rCCN3 in the same mice (left) and quantification thereof (right). Scale bars: 0.4mm. Data are presented as mean ± SEM; *n* = 3 biologically independent samples. Statistical significance was assessed using a two-tailed unpaired Student’s *t* test. Exact *P* values are indicated in the figure.

We next ablated Panx3⁺ cells in adult mice and again observed significant reductions in bone mass parameters (**Fig.5e,f & Extended Data Fig.5d**), demonstrating an ongoing requirement for this population in skeletal homeostasis. Given the intimate relationship between bone and hematopoiesis^31,40^, we further examined peripheral blood and bone marrow hematopoietic populations using established flow cytometric panels. Depletion of Panx3⁺ cells rapidly increased the frequency of circulating myeloid cells while reducing T cell abundance (**Extended Data Fig.5e,g**). In the bone marrow, this was accompanied by significant reductions in hematopoietic stem and progenitor populations in both frequency and absolute cell number (**Fig.5g & Extended Data Fig.5h**).

To directly assess SSC function, we compared the in vitro mineralization capacity of prospectively isolated Panx3⁺ and Panx3⁻ SSCs with SSCs isolated from Panx3-Cre^ERT2^/DTA mice. Panx3⁺ SSCs exhibited the greatest osteogenic potential, whereas Panx3⁻ and SSCs from DTA-treated mice showed significantly reduced Alizarin Red S staining (**Fig.5h**). These findings indicate reduced immediate skeletogenic activity among Panx3⁻ SSCs and following depletion of the Panx3⁺ compartment.. We next prospectively isolated SSCs from tamoxifen-treated 2-week-old Panx3-Cre^ERT2^/DTA mice and assessed their ossicle formation capacity by renal capsule transplantation into NSG recipients. Tamoxifen administration was continued in recipient mice during the first five days after transplantation. SSCs from DTA-treated mice generated significantly smaller grafts with reduced marrow cavity formation relative to equal numbers of control SSCs (**Fig.5i**). Further histological analysis revealed reduced or delayed endochondral ossification (Collagen-X) and a correspondingly smaller hematopoietic compartment, including fewer recruited myeloid cells (CD11b; **Extended Data Fig.5i**). Notably, Panx3 transcripts remained detectable in DTA SSC-derived grafts, suggesting that residual Panx3⁻ SSCs retained the capacity to generate Panx3⁺ SSCs after transplantation, albeit with delayed kinetics. These findings establish a functional requirement for the Panx3⁺ state in efficient skeletogenesis and formation of a hematopoiesis-supportive niche, while demonstrating that residual Panx3⁻ SSCs retain the capacity to acquire this state and regenerate skeletal tissue with delayed kinetics.

To more closely define the role of Panx3⁺ cells in skeletal regeneration, we then revisited our stabilized bi-cortical fracture model. Ablation of the Panx3-lineage one month before fracture significantly reduced trabecular bone volume within the callus relative to controls (**Extended Data Fig.5j**). Because pre-existing Panx3⁺ cells do not contribute directly to fracture healing (**Fig.4f**), this phenotype may instead reflect compromised skeletal integrity following lineage depletion. Supporting a direct requirement during repair, DTA-mediated ablation initiated at the time of fracture and maintained for the first five days of healing impaired callus formation and mineralization significantly (**Fig.5j**). As observed following renal capsule transplantation of SSCs, Panx3⁺ cell depletion impaired endochondral ossification and was accompanied by a failure to establish the hematopoietic compartment as part of callus remodeling, as evidenced by the absence of CD45 and cKit expression within callus tissue (**Extended Data Fig.5k**).

We next asked whether Panx3⁺ cells contribute to regeneration of AC following microfracture (MF), a setting in which we have shown SSCs are required for repair^21^. To determine whether injury recruits or induces Panx3⁺ cells within an otherwise Panx3⁻ AC environment, we introduced 1mm drill holes into the subchondral bone and implanted microgels containing PBS or recombinant CCN3 (rCCN3), a factor previously shown to promote SSC-mediated endochondral ossification even in aged settings^41^. MF alone predominantly generated fibrous tissue, whereas rCCN3 produced a proteoglycan-rich regenerate with features of hypertrophy by 4-weeks, as indicated by Safranin O staining (**Fig.5k**). Strikingly, DTA-mediated ablation of Panx3⁺ cells beginning at the time of MF drastically impaired rCCN3-induced de novo Aggrecan⁺ cartilage formation (**Fig.5k,l & Extended Data Fig.5l**). Lineage tracing in intact Panx3-Cre^ERT2^/Ai9 mice revealed extensive contribution of Panx3-lineage cells to newly formed subchondral bone during MF-induced remodeling, whereas hypertrophic cartilage at the joint interface was predominantly unlabeled (**Extended Data Fig.5m**). Thus, despite their absence from homeostatic AC, cells entering the Panx3⁺ state become functionally required for the regenerative response following injury. The unlabeled cartilage could arise from distinct local chondrogenic populations, including Procr⁺ AC progenitors^20^, or from cells that acquired the Panx3⁺ state after the initial 5-day tamoxifen-labeling window. Under rCCN3, however, Panx3⁺ cells labeled during the initial post-injury period contributed predominantly to remodeled subchondral bone rather than cartilage. These findings suggested that acquisition of the Panx3⁺ state does not itself specify regenerative fate and raised the possibility that subsequent lineage output remains instructible by the local niche.

### Spatial niche reconstruction reveals instructive control of Panx3-lineage fate

The preferential contribution of early injury-induced Panx3-lineage cells to subchondral bone under rCCN3 led us to investigate whether their regenerative fate could be redirected by alternative environmental signals. We therefore returned to the spatially resolved cartilage microexplant dataset generated in **Fig. 1** to identify signals distinguishing AC from GP and FX. Because newly established AC lacks appreciable Panx3⁺ SSC activity yet maintains a stable non-hypertrophic identity, we reasoned that its native niche might contain signals capable of directing injury-induced Panx3-lineage cells toward cartilage rather than endochondral progression. Mapping individual transcriptomes to cells within intact AC, GP and FX microexplants, this approach enabled direct interrogation of site- and zone-specific signaling programs (**Fig.1j & Extended Data Fig.1g**).

We applied LIANA to identify ligand-receptor interactions enriched in AC versus GP and FX populations, revealing distinct signaling features of AC (**Extended Data Fig.6a,b**)^42^. Specifically, the hyaline cartilage subcluster in our scRNAseq analysis (**Fig.1d,e & 6a,b, Extended Data Fig.6a,b**) was characterized by *Wif1*, *Lef1*, *Cytl1* and *Cxcl14* expression and enriched Wnt, Tgfβ, and Fgf signaling. The coexpression of *Wif1*, an extracellular Wnt antagonist, and *Lef1*, a downstream Wnt/β-catenin effector, suggested a Wnt regulated feedback state rather than simple pathway suppression. This could serve a mechanism that constrains excessive Wnt activity to maintain cartilage identity. LIANA further identified a predicted interaction between Wif1 and Ryk, an atypical Wnt receptor containing an extracellular WIF domain, suggesting an additional interface through which Wnt signaling could be modulated within the AC niche. Expression of *Fgfr1/2/3*, *Sdc4*, and *Thbs4* in AC further suggested responsiveness to extracellular matrix-associated growth factor and mechanical cues. *Fgf2* was among the relevant ligands expressed in hyaline cartilage, while its receptor *Fgfr3* was also present in the Panx3⁺ subpopulation. Spatial reconstruction of cartilage microexplants further showed *Wif1*- and *Fgfr3*-expressing cells along the superficial layer of AC (**Fig.6c & Extended Data Fig.6c**). Importantly, GP microexplants expressed Ryk and Fgfrs, indicating that GP-resident SSCs might possess the signaling machinery to respond to AC-associated Wnt and Fgf cues.

**Figure 6:**
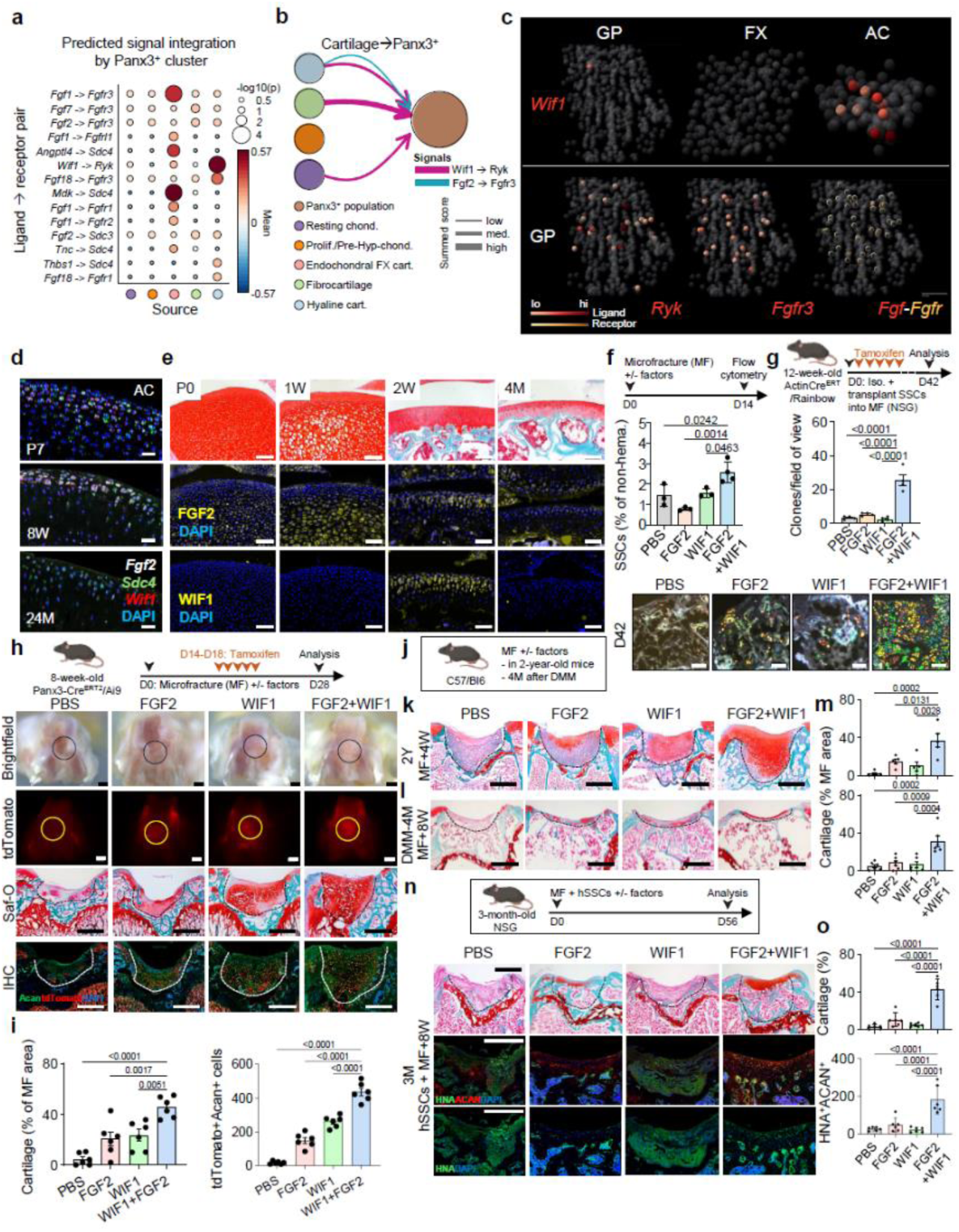
Native niche signals instruct the regenerative fate of Panx3-lineage cells. **(a)** LIANA-based ligand-receptor interaction analysis based on scRNAseq data of cartilage microexplants for selected significant ligand-receptor interactions of subpopulations. **(b)** Distinct signal integration of Panx3^+^ cluster based on CellChat. **(c)** Single cell spatial reconstruction of cartilage microexplants showing *Wif1* gene expression in individual cells across cartilage sites (top) and receptors (*Ryk*, *Fgfr3*) and ligand-receptor (*Fgf-Fgfr*) in GP (bottom). **(d)** RNAScope images of articular cartilage (AC) area at different postnatal ages. Scale bars: 50um. **(e)** AC area staining in P0, 1-, 2-, and 4-week-old mice showing Safranin O (top), IHC for Fgf2 (middle) and IHC for Wif1 (bottom). Scale bars: 100um. **(f)** Flow cytometric analysis of MF regenerate tissue for phenotypic SSCs in different treatment groups. Data are presented as mean ± SEM; *n* = 3-4 biologically independent samples. Statistical significance was assessed by one-way ANOVA followed by Tukey’s multiple-comparisons test. Exact *P* values for the indicated comparisons are shown in the figure. **(g)** MF experiment in 12-week-old Actin-Cre^ERT^/Rainbow mice with quantification of clones (top) and representative fluorescence images of regenerate area (bottom).Scale bars:100um. Data are presented as mean ± SEM; *n* = 4 biologically independent samples. Statistical significance was assessed by one-way ANOVA followed by Tukey’s multiple-comparisons test. Exact *P* values for the indicated comparisons are shown in the figure. **(h)** MF experiment in Panx3-Cre^ERT2^/Ai9 mice. Representative images showing photograph of joint area with focal defect (top), tdTomato signal (second from top), Safranin O staining of corresponding tissue sections (third from top) and IHC of tdTomato signal with Acan (bottom). Scale bars: 0.4mm. **(i)** Quantification of formed cartilage based on Safranin O staining (left) and tdTomato/Acan double-positive cells in regenerate area (right). Data are presented as mean ± SEM; *n* = 6 biologically independent samples. Statistical significance was assessed by one-way ANOVA followed by Tukey’s multiple-comparisons test. Exact *P* values for the indicated comparisons are shown in the figure. **(j)** Experimental outline for MF experiment with factors in aged 2-year-old mice and mice with 4-months established experimental OA via destabilization of the medial meniscus (DMM). **(k)** Representative Safranin O staining images of different groups in aged mice. Scale bars: 0.4mm. **(l)** Representative Safranin O staining images of different groups in DMM mice. Scale bars: 0.4mm. **(m)** Quantification of cartilage formed based on Safranin O stain in aged (top) and DMM mice (bottom). Data are presented as mean ± SEM; *n* = 5-6 biologically independent samples. Statistical significance was assessed by one-way ANOVA followed by Tukey’s multiple-comparisons test. Exact *P* values for the indicated comparisons are shown in the figure. **(n)** Experimental outline for MF experiment with factors and human SSCs in NSG mice (top). Representative Safranin O staining (top), scale bars: 0.4mm, IHC co-staining of Human Nuclear Antigen (HNA) with ACAN (middle) and HNA alone (bottom). **(o)** Quantification of cartilage formed (top) and human SSC ACAN+ cells (bottom). Data are presented as mean ± SEM; *n* = 6 biologically independent samples. Statistical significance was assessed by one-way ANOVA followed by Tukey’s multiple-comparisons test. Exact *P* values for the indicated comparisons are shown in the figure.

We therefore prioritized Wif1-mediated Wnt modulation together with Fgf2-Fgfr3 signaling as complementary candidate features of the hyaline cartilage niche. Panx3⁺ cells were absent from AC throughout development, indicating that these cues normally establish cartilage identity independently of Panx3⁺ SSCs. We reasoned that these AC-associated signals could instead be redeployed after injury to instruct cells entering the Panx3⁺ state toward stable cartilage while limiting hypertrophic and osteogenic progression. Longitudinal RNA labeling and antibody staining of joints further corroborated the distinct spatial and temporal expression patterns of these factors (**Fig.6d,e**). Fgf2 was abundant in developing AC but vanished upon skeletal maturation, whereas Wif1 showed transient expression in the joints of 1- and 8-week old mice at the RNA level and was detectable at the protein level specifically in 2-week-old AC. These transient patterns suggest complementary roles for Fgf2 and Wif1 in establishing hyaline cartilage identity during development. They also provided a rationale for leveraging developmental AC signals during adult repair.

We next tested whether these AC-specific signals could be leveraged to modulate SSC function. Prospectively isolated SSCs were stimulated with chondrogenic or osteogenic cocktails in the presence or absence of recombinant Wif1 and Fgf2 in vitro. Wif1 alone did not alter mineralization, whereas Fgf2, either alone or combined with WIF1, suppressed osteogenic potential (**Extended Data Fig.6d**). Conversely, Fgf2 increased proteoglycan production as measured by Alcian Blue staining, with a further increase upon combined Wif1 and Fgf2 treatment. We next tested these factors in the MF model, delivering recombinant Wif1 and Fgf2 locally using microgels placed at the time of MF. Flow cytometric analysis at day 14 revealed that combined Wif1 and Fgf2 treatment increased the frequency of SSCs and BCSPs at the injury site, suggesting enhanced progenitor recruitment with potential to support hyaline cartilage formation (**Fig.6f & Extended Data Fig.6e**). Similarly, transplantation of prospectively isolated Rainbow-reporter SSCs into MF sites and tamoxifen-pulsed upon delivery showed a marked increase in clonal activity only with combined Wif1 and Fgf2 treatment, whereas either factor alone had little effect (**Fig.6g & Extended Data Fig.6f**). Thus, coordinated Wif1 and Fgf2 signaling expanded SSC activity while shifting their functional output toward chondrogenesis and away from osteogenic differentiation.

To determine whether this niche environment could redirect the regenerative output of cells entering the Panx3⁺ state toward stable cartilage formation, we performed MF in Panx3-Cre^ERT2^/Ai9 mice with local delivery of Wif1 and Fgf2. At 4-weeks, the combined Wif1 and Fgf2 treatment produced significantly greater cartilage formation, as assessed by Safranin O and immunostaining, whereas PBS controls and single factor treatments showed features of fibrocartilage and hypertrophic differentiation, respectively (**Fig.6h,i & Extended Data Fig.6g**). Whereas Panx3⁺ cells labeled during rCCN3-mediated repair contributed predominantly to remodeled subchondral bone with little contribution to cartilage, combined Wif1 and Fgf2 treatment resulted in prominent Panx3-lineage contribution to Aggrecan⁺ cartilage (**Fig. 6h**). Thus, cells entering the Panx3⁺ state are not irreversibly committed to a predetermined regenerative trajectory. Instead, their subsequent lineage output remains malleable by the local signaling environment. Durable cartilage formation and native-like tissue remodeling with combined factors were similarly observed in wild-type mice at 4- and 8-weeks after MF (**Extended Data Fig.6h**).

To assess efficacy in clinically relevant settings, we extended these studies to aged mice and mice with experimental OA induced by destabilization of the medial meniscus (DMM). Strikingly, combined Wif1 and Fgf2 treatment consistently promoted hyaline-like cartilage regeneration in 1-year-old, 2-year-old and OA mice, which persisted up to 20-weeks (longest timepoint investigated) after extensive remodeling of the articular surface following MF (**Fig.6j-m & Extended Data Fig.6l,m**). Finally, to assess translational potential, we transplanted patient-derived human SSCs embedded in microgels with or without Wif1 and Fgf2 into MF sites of adult NSG mice. Primary human SSCs integrated into the joint and contributed to the regenerate, but only double factor-treated human SSCs consistently generated Aggrecan⁺ chondrocytes 8-weeks after MF (**Fig.6n,o**). Intriguingly, donor cells also contributed extensively to remodeled subchondral tissue, demonstrating retention of both chondrogenic and osteogenic potential. In sum, these findings demonstrate that acquisition of a regenerative stem cell state and determination of its subsequent lineage fate are separable processes. Native niche signals can be redeployed to redirect the output of injury-induced SSCs toward stable hyaline-like cartilage, including in aged, osteoarthritic and human settings.

## Discussion

Here, we identify Panx3 as a marker of a dynamic state within bona fide SSCs and establish a genetic model for interrogating this prospectively defined stem cell compartment in vivo. SSCs have traditionally been defined through combinatorial cell-surface markers and functional assays, enabling prospective isolation and characterization of their lineage hierarchy but limiting direct interrogation within their endogenous environment. By integrating spatial single cell transcriptomics, lineage mapping, prospective isolation, serial transplantation and ablation, we bridge this gap between functional definition and longitudinal interrogation in vivo. Panx3 selectively marked phenotypic SSCs with long-term clonal persistence, self-renewal and multipotent skeletal output, whereas Panx3⁻ phenotypic SSCs could give rise to Panx3⁺ SSCs. Thus, Panx3 does not define a fixed SSC identity but an acquirable functional state. Together with its engagement across developmental, systemic and regenerative settings, these findings suggest that anatomical origin and stem cell state represent separable dimensions of SSC identity.

The prominent localization of Panx3⁺ SSCs within the GP places this dynamic state within an increasingly heterogeneous stem cell compartment^18,43^. Postnatal Pthrp⁺ RZ cells generate columnar chondrocytes, osteoblasts and marrow stroma^10^, whereas Foxa2⁺ cells define long-term RZ stem cells^11^ and additional SSC populations extend across the GP-metaphyseal axis^44–46^. Panx3⁺ SSCs are enriched in the RZ-to-PHZ axis and exhibit distinct responses to cellular ablation, suggesting a functional contribution to GP organization that differs from previously characterized RZ populations. However, these differences do not establish mutually exclusive SSC identities and may instead reflect distinct functional states within interconnected progenitor populations. These findings implicate the Panx3⁺ state in progression towards hypertrophic differentiation, although whether Panx3 directly regulates this transition or marks cells undergoing it remains unclear. Prior studies placing Panx3 at the proliferative-to-prehypertrophic transition and early osteogenesis^22,29,47^, together with the absence of a major phenotype following osteoblast-restricted deletion^38^, support a function preceding terminal differentiation. Our findings connect these established Panx3 functions to SSC activity revealing their engagement within a functionally validated stem cell compartment.

This coexistence of stem cell function with early chondrogenic and prehypertrophic programs challenges a simple hierarchical view in which acquisition of differentiation-associated programs necessarily reflects progressive loss of stemness. Panx3⁺ SSCs retain long-term clonal activity and multipotent skeletal output, while Panx3⁻ SSCs can enter the Panx3⁺ state. Dynamic state acquisition is therefore compatible with durable stem cell behavior. The successive embryonic and neonatal waves of Panx3⁺ activity similarly do not imply that individual Panx3⁺ SSCs are intrinsically transient. Instead, they indicate that new SSCs can engage this state at different stages of skeletogenesis while cells already occupying it can persist and sustain clonal output. This distinction may help reconcile the growing number of anatomically and transcriptionally distinct skeletal progenitor populations with our finding that a common functional state can be acquired across otherwise distinct SSC populations.

The ability to genetically follow bona fide SSCs also provides a means to examine how this compartment responds to systemic signals relevant to skeletal aging and disease, e.g., osteoporosis. Panx3-lineage activity declined with age but persisted within adult growth plate and trabecular compartments. Depletion of Panx3⁺ cells reduced bone mass and impaired skeletal maintenance. Conversely, intermittent PTH1-34 increased Panx3⁺ clonal activity and enhanced subsequent lineage contribution across the skeleton. Although Panx3 has been associated with endochondral ossification, the expansion of Panx3⁺ clones along metaphyseal trabecular surfaces suggests that this state may also support direct osteogenic bone formation. These findings support Panx3 as a shared skeletogenic SSC state whose tissue-forming output depends on anatomical and environmental cues. Importantly, the response to PTH demonstrates that bona fide SSC activity remains pharmacologically responsive in the aged skeleton. Current osteoanabolic therapies do not selectively target SSCs. Their effects on this compartment have been difficult to interrogate genetically. Panx3-Cre^ERT2^ therefore provides an experimental framework for determining how SSC function changes with skeletal aging and osteoporosis, how existing anabolic therapies influence these cells and whether direct manipulation of SSC states can elicit distinct or complementary skeletal responses. The accompanying defects in hematopoietic stem and progenitor populations following Panx3⁺ cell depletion further connect SSC activity to formation and maintenance of marrow-supportive skeletal niches, consistent with established SSC functions^1,8^ and their coordinated decline with aging^31^.

A central implication of these findings is that a common stem cell-associated state can be engaged across anatomically distinct skeletal progenitor populations. Pre-existing Panx3-lineage cells did not contribute detectably to diaphyseal fracture repair, whereas labeling initiated at injury marked extensive chondrogenic and osteogenic callus populations. Because periosteal SSCs normally support intramembranous bone formation but acquire endochondral properties following fracture^12^, they represent one potential source of these injury-induced Panx3⁺ cells. Fgfr3⁺ osteochondrogenic progenitors spanning endosteal and periosteal compartments may provide another origin^15,48^. Therefore, the Panx3⁺ state can arise through at least two distinct modes. One, expansion of pre-existing Panx3⁺ SSCs in response to systemic anabolic stimulation and, two, local acquisition by skeletal progenitors following injury. Together with the successive developmental cohorts entering this state, these findings demonstrate that a common stem cell-associated program can be engaged by different environmental inputs without requiring a single anatomical cell source.

Just as acquisition of the Panx3⁺ state is separable from anatomical origin, its subsequent lineage output depends on the local environment. This distinction was particularly evident during AC regeneration. Although Panx3⁺ SSCs were absent from homeostatic AC, which contains Procr⁺ progenitors supporting cartilage maintenance^20^, Panx3⁺ cells became functionally important following MF. Under rCCN3 treatment, cells labeled during the early injury response contributed predominantly to remodeled subchondral bone, whereas combined Wif1 and Fgf2 redirected Panx3-lineage contribution towards Aggrecan⁺ cartilage. Guided by three-dimensional single cell reconstruction of native cartilage niches, this combination enhanced SSC recruitment and clonal expansion, promoted chondrogenesis and restrained hypertrophic and osteogenic progression. These effects were not reproduced by either factor alone. Co-expression of Wif1 and Lef1 suggests feedback regulation of Wnt activity, complemented by FGF signaling within the native AC niche. Thus, spatially defined niche signals can redirect the regenerative output of injury-responsive SSCs without predetermining their fate upon acquisition of the Panx3⁺ state. The resulting cartilage persisted in aged and osteoarthritic joints, while analogous treatment supported cartilage formation by human SSCs, highlighting the potential to harness endogenous niche programs for regeneration across age, disease and species.

The environmental responsiveness of Panx3⁺ SSCs parallels our previous finding that prospectively defined human SSC subtypes can interconvert in response to environmental cues^3^, suggesting that dynamic functional states may be a conserved feature of SSC biology. Related plasticity among periosteal and craniofacial progenitors further supports a view in which developmental origin and anatomical location shape, but do not irreversibly determine, SSC behavior. Altogether, our results challenge rigid hierarchical models of stem cell identity and instead distinguish anatomical origin, functional state and subsequent lineage fate as interconnected but separable dimensions. Panx3 provides genetic access to this dynamic relationship, while three-dimensional reconstruction of native niches reveals how environmental signals can shape its regenerative output. This framework offers new opportunities to investigate how SSC plasticity is maintained, altered with aging and disease or harnessed therapeutically to restore skeletal tissues.

## Methods

### Mouse strains

All mouse experiments complied with relevant ethical regulations and were conducted under protocols approved by the Stanford University Administrative Panel on Laboratory Animal Care and the University of California, Davis Institutional Animal Care and Use Committee (IACUC). Mice were housed in research animal facilities at Stanford University and UC Davis in accordance with institutional guidelines. Animals had ad libitum access to food and water and were maintained in temperature- and humidity-controlled microisolator cages under a 12-h light/12-h dark cycle.Unless otherwise specified, experiments were performed using male and female C57BL/6J mice (The Jackson Laboratory, JAX: 000664) at the indicated ages. Rainbow barcoding and lineage-tracing experiments were performed using young and aged (specific ages indicated) male Actin-CreERT2;R26VT2/GK3 or Panx3-CreERT2;R26VT2/GK3 mice bred and maintained in our laboratory. Panx3-CreERT2 mice were crossed with Ai9 reporter mice (B6.Cg-Gt(ROSA)26Sor^tm9(CAG-tdTomato)Hze/J; The Jackson Laboratory, JAX: 007909) or DTA mice (B6.129P2-Gt(ROSA)26Sor^tm1(DTA)Lky/J; The Jackson Laboratory, JAX: 009669) to generate the corresponding Panx3-CreERT2;Ai9 and Panx3-CreERT2;DTA strains. Appropriate offspring were subsequently intercrossed as required to obtain the experimental genotypes used in this study.

### Human primary cells

Human skeletal stem cells (hSSCs) were isolated from fracture callus tissues collected 3-5 days after injury during open reduction and internal fixation procedures. Tissue procurement and processing were performed in accordance with protocols approved by the Stanford University Institutional Review Board (IRB-35711). Informed consent was not required because the collected specimens were classified as discarded biological waste. No restrictions were applied based on donor race, sex, or age. Immediately after excision, specimens were placed on ice and processed for hSSC isolation as described below.

### Tamoxifen treatment

Tamoxifen (Sigma, T5648) was dissolved in corn oil at a concentration of 20 mg/ml and administered by oral gavage at 200 mg/kg body weight at the indicated time points. For neonatal pups, 3 μl of tamoxifen solution (40 mg/ml in corn oil) was administered orally.

### Genotyping

Mouse tail-tip genomic DNA was prepared using the Mouse Quick Genotyping DNA Preparation Kit (Bioland Scientific LLC, GT01-01) according to the manufacturer’s instructions. Genotyping PCR was performed using GoTaq® G2 Hot Start Master Mix (Promega, M7422). Primer sequences are provided in key resourse table.

### Flow cytometric analysis and cell isolation

Mouse SSC lineage populations were isolated using previously described cell-surface marker profilesgulat^1,49^. In brief, femurs were dissected, cleared of surrounding soft tissue, and finely minced using a sterile razor blade. The resulting tissue fragments were digested in M199 medium (Thermo Fisher Scientific, Cat# 11150067) containing 2.2 mg ml⁻¹ collagenase II (Sigma-Aldrich, Cat# C6885) at 37°C for 60 min. Dissociated cells were passed through a 70-μm nylon filter, washed in staining medium consisting of 10% fetal bovine serum (FBS) in PBS, and pelleted at 200 × g at 4°C. The cell pellet was resuspended in staining medium and subjected to ACK lysis for 5 min to remove erythrocytes. Cells were then washed again in staining medium, pelleted at 200 × g at 4°C, and stained with fluorochrome-conjugated antibodies against CD45 (1:200, BioLegend, Cat# 103110), TER119 (1:200, BioLegend, Cat# 116210), CD51 (1:50, BioLegend, Cat# 104106,104104), TIE2 (1:20, BioLegend, Cat# 124002), THY1.1 (1:100, BioLegend, Cat# 202520), THY1.2 (1:100, BioLegend, Cat# 140332), 6C3 (1:100, BioLegend, Cat# 108312), CD200 (1:100, BD OptiBuild, Cat# 745402), and CD105 (1:50, BioLegend, Cat# 120410), together with BV785-conjugated streptavidin(1:100, BioLegend, Cat# 405249), before flow cytometric analysis and sorting. Human SSCs were isolated from fracture callus tissue using the same digestion and single-cell preparation procedures described above for mouse SSC isolation. The resulting cell suspensions were stained with fluorochrome-conjugated antibodies against PDPN (1:100, BioLegend, Cat# 337022), CD73 (1:50, BioLegend, Cat# 344016), CD164 (1:100, BioLegend, Cat# 324808), CD146 (1:100, BioLegend, Cat# 342101), CD31 (1:100, BioLegend, Cat# 303120), TIE2 (1:100, BioLegend, Cat# 334204), CD235a/b (1:100, BioLegend, Cat# 306612), and CD45 (1:100, BioLegend, Cat# 304022) together with APC/Cyanine7-conjugated streptavidin(1:100, BioLegend, Cat# 405208), followed by flow cytometric analysis and cell sorting. For analysis of bone marrow hematopoietic populations, bone marrow was collected from femurs by centrifugation at 12,000 × g for 1 min at 4°C. Red blood cells were depleted by ACK lysis, and the remaining cells were washed with FACS buffer before staining with antibodies against hematopoietic lineage markers(BioLegend, Cat#1 33310), CD127(BioLegend, Cat# 135035), CD117(BioLegend, Cat# 105826), Sca-1(BioLegend, Cat# 160904), CD16/32((BioLegend, Cat# 156606), CD34(BD OptiBuild, Cat# 742971), CD135(BioLegend, Cat# 135312), and CD150(BioLegend, Cat# 115929), each at a 1:100 dilution. For peripheral blood analysis, blood was collected by terminal cardiac puncture into EDTA-containing tubes. Samples were diluted with FACS buffer and pelleted at 200 × g at 4°C. Red blood cells were depleted by ACK lysis for 5 min, after which cells were washed twice with FACS buffer and stained with fluorochrome-conjugated antibodies against TER119(BioLegend, Cat# 116210), CD45(eBioscience, Cat# 11-0451-82), B220(BioLegend, Cat# 103224), CD11b(BioLegend, Cat# 101216), CD3(BioLegend, Cat# 100236), and GR-1(BioLegend, Cat# 108443), each at a 1:200 dilution. Flow cytometry was performed using a FACSAria II instrument (BD Biosciences) or an Aurora CS cell sorter (Cytek Biosciences). Gating strategies were established using fluorescence-minus-one controls. DAPI were added to the sample right before analysis to exclude nonviable cells. All cell populations were sorted for purity.

### *In vitro* cell culture

Cells were maintained in MEM-α supplemented with 10% fetal bovine serum (FBS, HyClone, Cat# SH30396.03) and 1% penicillin–streptomycin (Thermo Fisher Scientific, Cat# 15140-122). For in vitro skeletogenic differentiation assays, 12,000 FACS-purified mouse SSCs were cultured in individual wells of 24-well tissue culture plates. Cells were washed with PBS(Gibco, Cat# C14190500BT), trypsinized, and transferred to the indicated differentiation conditions. For osteogenic differentiation, cells were cultured for 14 days in MEM-α (Fisher Scientific, Cat# 12561-056) supplemented with 10% FBS, 100 nM dexamethasone(Sigma-Aldrich, Cat# D2915), 100 μg ml⁻¹ L-ascorbic acid-2-phosphate(Sigma-Aldrich, Cat# A8960), and 10 mM β-glycerophosphate(Sigma-Aldrich, Cat# G9891). For chondrogenic differentiation, micromass cultures were generated by placing a 5-μl droplet of cell suspension at approximately 1.5 × 10⁷ cells ml⁻¹ in the center of each well of a 48-well plate. Cells were incubated for 2 h at 37°C to allow attachment before the addition of pre-warmed chondrogenic medium consisting of high-glucose DMEM(Gibco, Cat# 11965092) supplemented with 10% FBS, 100 nM dexamethasone(Sigma-Aldrich, Cat# D2915), 1 μM L-ascorbic acid-2-phosphate(Sigma-Aldrich, Cat# A8960), and 10 ng ml⁻¹ TGF-β1(R&D, Cat# 7666-MB). Where indicated, recombinant proteins including WIF-1 (R&D, Cat# 135-WF), FGF2 (R&D, Cat# 3139-FB), and syndecan-4 (R&D, Cat# 6267-SD) were added to the differentiation medium. Chondrogenic cultures were maintained for 21 days. At the end of differentiation, cultures were washed with PBS and fixed with 4% paraformaldehyde (PFA). Osteogenic differentiation was assessed by staining with Alizarin Red S (Sigma-Aldrich, Cat# A5533-25G), whereas chondrogenic differentiation was assessed by Alcian Blue staining (Sigma-Aldrich, Cat# A3157). Following imaging using a flatbed scanner, bound Alizarin Red S was extracted with 10% acetic acid and quantified by measuring absorbance at 450 nm. Alcian Blue was extracted with 12 M guanidine(Sigma-Aldrich, Cat# G4505) in 1 M HCl and quantified by measuring absorbance at 630 nm.

### Subcapsular renal transplantation

Subcapsular renal transplantation was performed as previously described^9,49^.Briefly, NSG mice were anesthetized with aerosolized isoflurane and analgesia was administered before surgery, and the kidney was exposed through a 10-mm dosal incision. A suspenstion containing 5000 FACS-purified mouse SSC in 5 μl of Matrigel(Corning, Cat# 354230) together with PBS or recombinant proteins including WIF-1 (R&D, Cat# 135-WF), FGF2 (R&D, Cat# 3139-FB), and syndecan-4 (R&D, Cat# 6267-SD) was allowed to polymerize at room temperature for 10min and then transplanted beneath the renal capsule. The kidney was returned to the peritoneal cavity and the incision was closed with sutures and surgical staples. Grafts were harvested at indicated timepoints for histological analysis, FACS analysis and/or secondary transplantation. Secondary transplantations were performed using the same procedure.

### Bi-cortical fracture model

Adult male mice of the indicated genotypes were anesthetized with aerosolized isoflurane, and analgesia was administered prior to surgery. The distal femur was exposed through lateral dislocation of the patella. A 25-gauge needle (BD Biosciences) was inserted through the femoral condyles to provide intramedullary fixation, and a transverse mid-diaphyseal femoral fracture was generated using fine surgical scissors. The patella was then repositioned, and the muscle and skin incisions were closed with 6-0 nylon sutures (Ethicon). Mice were sacrificed at the indicated time points after fracture for further analysis.

### Mouse destabilization of the medial meniscus (DMM) model

DMM surgery was performed as previously described^50^. Briefly, adult male mice of the indicated genotypes were anesthetized with aerosolized isoflurane, and the medial meniscotibial ligament was transected through a medial parapatellar approach to destabilize the medial meniscus. The joint capsule and skin were closed with 6-0 nylon sutures (Ethicon). Mice were maintained for 4 months after surgery to allow osteoarthritis development before used for subsequent experimentation.

### Mouse microfracture model

Microfracture surgery was performed as previously described^21^. Briefly, adult mice of the indicated genotypes were anesthetized with aerosolized isoflurane. Following lateral patellar dislocation, a full-thickness osteochondral defect was generated in the trochlear groove of the distal femur using a 0.8-mm dental drill, penetrating the subchondral bone to establish communication with the marrow cavity. Recombinant proteins including NOV/CCN3 (R&D, Cat# 1976-NV), WIF-1 (R&D, Cat# 135-WF), FGF2 (R&D, Cat# 3139-FB), and syndecan-4 (R&D, Cat# 6267-SD) at the indicated concentrations were encapsulated in 1% MVG sodium alginate(PronovaNovamatrix, Cat# 4200101) and polymerized within molds generated using a 1-mm biopsy punch in 1.5% agarose containing 1 M CaCl₂. The alginate constructs were allowed to solidify on ice for 30 min before implantation into the defect. For human SSC transplantation experiments, 25,000 human SSCs suspended in 1 μl of Matrigel were applied directly to the defect prior to implantation of the alginate construct. The patella was then repositioned, and the joint capsule and skin were closed with 6-0 nylon sutures (Ethicon). Mice were harvested at the indicated time points for downstream analyses.

### Parathyroid hormone (hPTH 1-34) treatment

Human parathyroid hormone 1–34 (hPTH(1–34); Sigma, P3796) was dissolved in PBS at a concentration of 2.5 μg/μl as a stock solution and diluted to 0.1 μg/μl before use. Mice received subcutaneous injections of hPTH(1–34) at a dose of 40 μg/kg body weight, administered 5 times per week for 2-4 consecutive weeks depending on the experiment as indicated.

### Click-iT™ EdU cell proliferation assay

EdU (5-ethynyl-2′-deoxyuridine) was dissolved in DMSO at 100 mg/ml as a stock solution and diluted in PBS to a final concentration of 20 mg/ml immediately before use. For pulse-labeling experiments, mice received a single intraperitoneal (i.p.) injection of EdU (50 mg/kg body weight) and were harvested at the indicated time points. For label-retention experiments, mice received EdU (50 mg/kg body weight, i.p.) once daily for 5 consecutive days. EdU incorporation was detected using the Click-iT™ EdU Imaging Kit with Alexa Fluor™ 488 azide (Thermo Fisher Scientific, C10086) according to the manufacturer’s instructions. Fluorescent images were acquired by confocal microscopy.

### Histology and immunofluorescence staining

Freshly isolated mouse tissues were fixed in 4% paraformaldehyde (PFA, Thermo Fisher Scientific, Cat# 047340.9L) at 4 °C for 24 h and decalcified in 400 mM EDTA(Invitrogen, Cat# 15573-038) in PBS for 2 weeks. Tissues were subsequently cryoprotected in 30% sucrose(Fisher Scientific, Cat# S5-500), embedded in OCT compound, and sectioned at a thickness of 10 μm. Representative sections were stained with freshly prepared haematoxylin and eosin, Movat’s pentachrome,, safraninO and fastgreen, or TRAP. For immunofluorescence staining, tissue sections were permeabilized and blocked for 1 h at room temperature in blocking buffer consisting of 3% bovine serum albumin (BSA, Sigma-Aldrich, Cat# A2153), 1% Triton X-100(FisherScientific, Cat# AAA16046AP), and 0.5% Tween-20(Sigma-Aldrich, Cat# P7949) in PBS. Sections were then incubated with blocking or primary antibodies including goat F(ab) anti-mouse IgG H&L (Abcam, Cat# ab6668), mouse anti-human nuclear antigen (Abcam, Cat# ab191181), rabbit anti-tdTomato (CST, Cat# 20163), rabbit anti-aggrecan (Proteintech, Cat# 13880-1-AP), rabbit anti-COL1A1 (CST, Cat# 72026), rabbit anti-collagen X (Abcam, Cat# ab58632), rabbit anti-collagen X (Invitrogen, Cat# PA5-115039), rabbit anti-osteopontin (Proteintech, Cat# 22952-1-AP), rabbit anti-WIF1 (Invitrogen, Cat# PA5-76731), rabbit anti-FGF2 (Invitrogen, Cat# PA5-116495), rabbit anti-CXCL12/SDF-1 (Proteintech, Cat# 17402-1-AP), and rabbit anti-perilipin-1 (CST, Cat# 9349L), Alexa Fluor 488–conjugated anti-mouse/human CD11b (BioLegend, Cat# 101217), anti-mouse CD45 (BioLegend, Cat# 160306), and anti-mouse CD117 (BioLegend, Cat# 105816) overnight at 4 °C. Following primary antibody incubation, sections were washed three times for 15 min each in PBS containing 0.5% Tween-20 and subsequently incubated with DAPI and the appropriate fluorophore-conjugated secondary antibodies including Alexa Fluor 488–conjugated goat anti-rabbit IgG (Invitrogen, Cat# A11008), Alexa Fluor 647–conjugated goat anti-mouse IgG (Invitrogen, Cat# A21235), Alexa Fluor 488–conjugated goat anti-rat IgG (Invitrogen, Cat# A11006), and Alexa Fluor 750–conjugated goat anti-rabbit IgG (Invitrogen, Cat# A21039). for 1 h at room temperature. Sections were washed three additional times in PBS containing 0.5% Tween-20 before mounting with ProLong™ Glass Antifade Mountant (Thermo Fisher Scientific, P36984). Fluorescent images were acquired using confocal microscopy.

### RNA in situ hybridization

Freshly isolated mouse long bones were fixed overnight and decalcified in 400 mM ethylenediaminetetraacetic acid (EDTA; Invitrogen, Cat# 15573-038) in PBS (pH 7.2) at 4°C for 2 weeks. Samples were subsequently cryopreserved and sectioned at 10 μm thickness. RNA in situ hybridization was performed using the RNAscope® Multiplex Fluorescent Reagent Kit v2 (Advanced Cell Diagnostics, Cat# 323100) together with probes targeting mouse *Panx3* (Advanced Cell Diagnostics, Cat# 847551), *Fgf2* (Advanced Cell Diagnostics, Cat# 316851), *Wif1* (C2; Advanced Cell Diagnostics, Cat# 412361-C2), and *Sdc4* (C3; Advanced Cell Diagnostics, Cat# 473591-C3) according to the manufacturer’s instructions. Sections were hybridized with indicated probes. Following RNAscope detection, immunofluorescence staining was performed using indicated antibody, followed by a Alexa Fluor 750 conjugated secondary antibody. Nuclei were counterstained with DAPI.

### MicroCT analysis

Soft tissue and pin-free femurs were scanned after 24h of fixation using a Bruker SkyScan 1276 system (Bruker Preclinical Imaging) at a source voltage of 85 kV and current of 200 μA, with a 1-mm Al filter, a voxel size of 12 μm, and an image resolution of 2016 × 1344 pixels. Manufacturer-provided phantom standards were used for instrument calibration. Reconstructed datasets were analyzed using CTAn v1.17.7.2 and CTvox v3.3.0 software (Bruker).For analysis of trabecular bone in uninjured femurs, a region of interest comprising 200 consecutive sections was defined beginning 50 sections distal to the end of the growth plate. For fracture callus analysis, 150 sections on either side of the middle of the fracture gap were selected, yielding a total analysis region of 300 sections. The fracture callus located outside the cortical bone boundaries was then manually delineated in CTAn for quantitative analysis.

### Three-dimensional spatial single cell reconstruction of cartilage microexplants

For articular cartilage and growth plate cartilage samples, Actin-CreERT2;R26VT2/GK3 pups were administered 0.12 mg tamoxifen orally at P3 and tissues were harvested at approximately P14 to achieve maximal fluorescent color diversity. For fracture callus samples, adult Actin-CreERT2;R26VT2/GK3 mice received tamoxifen at 200 mg kg⁻¹ 1 month before bicortical fracture surgery and again immediately after surgery. Fracture calluses were harvested 8-9 days after injury, a time point selected to maximize Rainbow color diversity and cartilage abundance within the regenerating tissue. In total, eight microexplants with a total of 1,125 cells (growth plate 384; articular cartilage: 144; fracture callus: 597) were analyzed for single cell sequencing in this study. Freshly isolated and microdissected cartilage from tamoxifen induced Actin-CreERT2;R26VT2/GK3 mice was sectioned at 70um using a vibratome and evenly dissected into cartilage fragments by pressing through a metal mesh to generate microexplants containing a few hundred cells. Individual cartilage fragments were carefully transferred to an imaging chamber containing FACS buffer supplemented with ProLong™ Live Antifade Reagent (Invitrogen, P36975) and imaged by confocal microscopy (Zeiss, LSM 980 Plus) from both forward and reverse orientations to acquire z-stacks for spatial localization and fluorophore barcode identification of individual cells. Following imaging, the cartilage fragment was gently transferred to a low-protein-binding 1.7-ml microcentrifuge tubes containing 200 μl of digestion buffer and dissociated into single cells by incubation at 37 °C with agitation (500 rpm) for 30 min in a thermomixer. Digestion was terminated by the addition of 1 ml of FACS buffer, and samples were gently triturated for 5 min before fluorescence-activated cell sorting (FACS) with index sorting for plate-based single-cell RNA sequencing (Smart-seq2). Fluorophore intensities of individual cells were quantified independently from both confocal imaging and FACS data and integrated using the bioinformatic matching pipeline described below. Matched cells were subsequently linked to their spatial coordinates and transcriptomic profiles to reconstruct spatially resolved three-dimensional single-cell maps and infer cell-cell interactions.

### Plate-based Smart-seq2 single-cell RNA sequencing

Single-cell suspensions derived from target tissues were prepared for flow cytometric isolation as described above and previously published^3,9,31^. Single-cell RNA sequencing was performed using a plate-based Smart-seq2 workflow with minor modifications. Briefly, individual cells were index-sorted into 96-well plates containing 4 μl of lysis buffer composed of 1 U μl⁻¹ RNase inhibitor (Clontech, Cat# 2313B), 0.1% Triton (Thermo Fisher Scientific, Cat# 85111), 2.5 mM dNTPs (Invitrogen, Cat# 10297-018), 2.5 μM oligo-dT30VN (IDT, custom: 5′-AAGCAGTGGTATCAACGCAGAGTACT30VN-3′), and ERCC ExFold RNA Spike-In Mix 2 diluted 1:600,000 (Invitrogen, Cat# 4456739) in nuclease-free water (Thermo Fisher Scientific, Cat# 10977023). Immediately after sorting, plates were centrifuged at 3,000 × g for 30 s at 4°C, snap-frozen on dry ice, and stored at −80°C until cDNA synthesis. For reverse transcription and cDNA pre-amplification, lysis plates were thawed on ice and incubated at 72°C for 3 min, followed by immediate cooling on ice to promote annealing of the oligo-dT30VN primer. First-strand cDNA synthesis was performed using SMARTScribe reverse transcriptase (Clontech, Cat# 639538) together with a locked nucleic acid-containing template-switching oligonucleotide (TSO; Exiqon, custom: 5′-AAGCAGTGGTATCAACGCAGAGTACATrGrG+G-3′). cDNA was subsequently amplified using KAPA HiFi HotStart ReadyMix (Kapa Biosystems, Cat# KK2602) and IS PCR primers (IDT, custom: 5′-AAGCAGTGGTATCAACGCAGAGT-3′). To remove residual reaction components and oligonucleotides smaller than 400 bp, amplified cDNA was purified using 0.65– 0.75× volumes of calibrated AMPure XP beads (Beckman Coulter, Cat# A63882). The concentration and size distribution of cDNA from each well were assessed using a high-sensitivity AATI 96-capillary fragment analyzer with the HS NGS Fragment Kit (1–6,000 bp; Agilent). cDNA from individual wells was normalized to approximately 0.15 ng μl⁻¹ by dilution with EB buffer. The contents of four 96-well plates were then consolidated into 384-well plates without cherry-picking or removal of wells lacking detectable single-cell cDNA. Sequencing libraries were prepared from the 384-well plates using the Nextera XT kit (Illumina, Cat# FC-131-1096) in a semi-automated workflow. Barcoded libraries from individual wells were pooled, purified, and size-selected using sequential 0.35× and 0.75× Agencourt AMPure XP bead cleanups according to the Nextera XT protocol. Final library fragment size distributions and concentrations were assessed using a high-sensitivity AATI fragment analyzer. Pooled libraries were sequenced on an Illumina NovaSeq 6000 S4 flow cell to generate approximately 1–2 million 2 × 150-bp paired-end reads per cell.

### Single cell RNA-seq data processing

Single cell RNA-seq data were processed as previously described^3,9,31^. ScRNA-seq data were demultiplexed using bcl2fastq2 2.18 (Illumina). Raw reads were further processed using skewer v.0.2.2 for 3’ quality trimming, 3’ adaptor trimming, and removal of degenerate reads. Trimmed reads were then mapped to the mouse genome v.M20 using STAR v.2.6.1plus (with an average of more than 70% of uniquely mapped reads), and counts per million (CPM) was calculated using RSEM v.1.3.1plus. Data were explored and plots were generated using the Scanpy package (v.1.8.0.). To select for high-quality single cells, cells with fewer than 300 genes and fewer than 25,000 counts were excluded. Genes that were detectable in fewer than 3 cells across all cells were also removed. In addition, any cells exceeding 20% mitochondrial and 25% ribosomal gene content as well as with a ERCC fraction higher than 30% were excluded. After these stringent quality filtering steps 546 cells were used for downstream analysis (growth plate 257; articular cartilage: 44; fracture callus: 245). Raw CPM values were mean- and log-normalized. Batch correction using Scanpy integrated ComBat was applied for cells derived from separate 96-well processing plates followed by data scaling. Clustering of single-cell data was done using UMAP(0.5.7) and leidenalg (0.10.2). Principal component (PC) elbow plots were used to select the number of PCs for each clustering analysis. Genes were ranked and differential expression between groups was assessed by Wilcoxon rank-sum test (Mann–Whitney *U*) for identification of cluster-specific genes. Differences in developmental state were assessed with stemness score. To that end, we combined two complementary metrics: a CytoTRACE-like score reflecting gene expression complexity smoothed across nearest neighbors, and the log-transformed number of expressed genes per cell as a direct measure of gene complexity. Both metrics were standardized across all cells using z-scores, and the per-cell stemness score was computed as the mean of these two z-scores. Higher stemness scores indicate cells with greater transcriptional diversity and predicted early progenitor states, whereas lower scores correspond to more differentiated cell states. Cluster-level stemness scores were obtained by averaging per-cell scores within each leiden_final cluster, allowing ranking of clusters from most stem-like to most differentiated. To facilitate interpretation and visualization, per-cell stemness scores were then min-max normalized to a 0-1 range, where 1 represents the most stem-like/early progenitor cells and 0 the most differentiated cells. Cluster-level stemness scores were obtained by averaging normalized per-cell scores within each Leiden cluster, allowing ranking of clusters from most stem-like to most differentiated.Differentially expressed gene within each leiden cluster were tested for GO enrichment using the Enrichr^53^ Webserver version of GO Biological Processes. Only significantly enriched GO terms were considered (adj. *P* < 0.1) and provided combined statistical scores are shown in the figures.

### Three-dimensional reconstruction and matching of single cell RNA-sequencing and confocal data

Cartilage explants representing growth plate (GP), articular cartilage (AC), and fracture callus (FX) regions underwent Forward and Reverse confocal imaging to neutralize depth attenuation, assess segmentation reliability, and measure baseline noise. Cells were segmented via Cellpose 4.1.1 using tissue-specific voxel geometries extracted from CZI metadata, generating initial segmented confocal pools of cells. Scans were aligned via a rigid transformation (z/x flips, z-axis rotation, and translation) derived from phase cross-correlation on band-passed data (3–30 µm) to anchor registration to cellular architecture. Spatial transforms strictly required a +1 determinant. Median Intersection-over-Union (IoU) scores were 0.34 (AC), 0.28 (FX), and 0.21 (GP), representing 4.0–5.5 µm spatial offsets. This metric conservatively understates alignment quality by conflating registration error with intrinsic segmentation variance. Median nearest-neighbor distances (8.2–12.5 µm) generally exceeded the alignment offset, ensuring counterpart matching. In dense GP regions (<4.9 µm spacing), potential mispairings inject noise rather than false biological structure and are partly self-filtering: a mispaired cell yields a broad posterior and low confidence, making it more likely to fail the inclusion criteria. Rigid, rather than deformable, registration was maintained to preserve the independence of the two optical measurements. For a cell at depth *d* within a specimen of thickness *D*, the Forward and Reverse scans traverse complementary path lengths, so their geometric mean 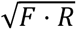 is independent of depth under Beer– Lambert attenuation. The cancellation follows from the acquisition geometry and requires no fitted attenuation coefficient, so its validity does not rest on a regression fit. All per-cell intensities used downstream are this geometric mean. Index-sorted FACS events were retroactively linked to spatial confocal coordinates. FACS intensities were quantile-normalized to confocal distributions, converted to relative color ratios, spatially detrended and paired with a fifth feature matching FACS FSC-A to confocal cellular volume. A Gaussian log-likelihood matrix incorporating heteroscedastic noise was resolved via linear-sum assignment (Hungarian algorithm). Robustness was bootstrapped 300 times utilizing the empirical variance between Forward and Reverse scans, producing initial index-sorted assignments of 290 for GP, 58 for AC, and 101 for FX. Assignments were preserved if confidence exceeded random chance (1/n) and IoU > 0. Passing matches beat random probability by 8–29× across datasets. Median posterior entropy spanned 73–92% of maximum, with the assigned cell accounting for 3–15% of posterior probability mass. The supported claim is therefore that each accepted match surpassed random chance by the specified margin with a verified dual-mask correspondence, rather than that deterministic uniqueness was achieved for every cell. The reconstruction is interpreted as an ensemble, its validity resting on the population-level tests below rather than on any individual pairing. This dual-criteria filtration yielded 247 valid assignments for GP (85% passage rate), 51 for AC (88%), and 95 for FX (94%). Finally, integrating these valid assignments with the SmartSeq2 expression matrix produced final spatially mapped cohorts of 60% for GP, 22% for AC, and 44% for FX. SmartSeq2 profiling evaluated selected transcripts as well as niche signaling pathways factors (*Wif1, Fgf2, Sdc4*), receptors (*Pth1r, Ryk, Fgfr1, Fgfr2, Fgfr3*), and Fgf-related factors (*Fgf1, Fgf7, Fgf18, Sdc3, Gpc4, Itgb3*). Composite pathway metrics were aggregated via predefined lists. Transcripts were projected into 3D models using absolute raw counts without z-scoring to preserve non-detection boundaries. Cellular volume expansion independently oriented the GP developmental vector at 164.5° (Spearman rho = +0.26) without transcriptomic input. Disabling transcriptomic spatial priors yielded a 95% identical assignment matrix. In this unbiased reconstruction, cell populations stratified into expected anatomical sequences (Jonckheere– Terpstra p=0.002), with Apoe⁺ at 0.44, Epyc⁺ at 0.51, and Panx3⁺ at 0.67. This spatial ordering persisted even when cell volume was neutralized as a matching feature (p=0.006). *Panx3* abundance significantly correlated with proximity to the pre-hypertrophic pole (Spearman rho = +0.19, p=0.0068). Median whole-transcriptome depth was 923 genes per cell for GP, 792 for AC and 1365 for FX. *Panx3* was detected in 33% of growth plate cells despite growth plate being shallower than callus. Across full datasets, *Wif1, Col1a1, Acan*, and *Panx3* exhibited significant site-dependent variation (p<0.05 post-Bonferroni correction). *Wif1* was found in 72% of AC and virtually absent elsewhere. Because spatial mapping evaluates intra-group topological organization, spatial figures strictly visualize structure. Full-dataset sequencing is utilized to validate absolute transcript prevalence or absence.

### Quantification and statistical analysis

Statistical significance between two groups was assessed using a two-tailed, unpaired Student’s t-test. For comparisons involving multiple groups, one-way ANOVA followed by either Turkey-Kramer’s multiple-comparisons test (for comparisons between 2 group) or Dunnett’s multiple-comparisons test (for comparisons to a control group) was performed, unless otherwise specified on the figure legends (GraphPad Prism, version 10). Statistical significance was defined as *p* < 0.05. All data points represent biological replicates and are presented as mean ± standard error of the mean (SEM), unless otherwise indicated in the figure legends.

### Data Availability

Generated scRNAseq data has been submitted and is available from GEO upon request. No original code was generated for this manuscript.

## Acknowledgments

We are grateful for valuable scientific discussions, feedback and laboratory management support from colleagues within UC Davis Department of Orthopaedic Surgery as well as the Chan and Weissman laboratories at Stanford University. We are grateful for technical assistance from Charlene Wang (Stanford) and Naymalis La Santa (UC Davis). We thank Stanford University Institute for Stem Cell Biology and Regenerative Medicine FACS core for experimental support, and NIH S10 1S10 OD028493 (PI: C.K.F.C). MicroCT work on the Bruker Skyscan 1276 was supported by NIH S10 Shared Instrumentation Grant (S10 OD023497, PI: Timothy C. Doyle). Funding for this work was provided by NIH-NIA K99/R00AG066963, NIA-NIA R01AG095155 to T.H.A. as well as W.M. Keck Foundation Bridge Funding Initiative support to T.H.A. and K.C.W., NIH-NIA K99/R00AG049958, Stanford Wu Tsai Human Performance Alliance Funds, Siebel Foundation, the Heritage Medical Foundation, Prostate Cancer Foundation, the American Federation for Aging Research (AFAR), Arthritis National Research Foundation (ANRF), a seed grant from the WHSDM Stanford Women’s Health and Sex Differences in Medicine Center and an endowment from the DiGenova Family to C.K.F.C. D.M. was supported by a National Institute of Arthritis and Musculoskeletal and Skin Diseases funded T32 training program in Musculoskeletal Health Research (T32 AR079099).

## Author contributions

Conceptualization and experimental design, Y.W., C.K.F.C. and T.H.A.; the single spatial transcriptomic approach was conceived by C.K.F.C.; the original mouse lines were generated by J.X. and G.M-A; methodology and data acquisition, Y.W., L.Z., K.C., E.H., J.S., K.D., H.M.S., N.R.M., K.C.W., E.C., D.M., M.Y.H., C.M., H.C., S.W. and W.L.; experimental analysis and data interpretation support, J.X., F.G., M.T.L., and I.L.W.; manuscript writing, review and editing, Y.W., L.Z. and T.H.A.; project supervision, M.T.L., I.L.W., C.K.F.C. and T.H.A.

## Extended Data Figure Legends

**Extended Data Figure 1: Pannexin-3 across skeletal tissues and its connection to mouse SSCs. (a)** Dot plot showing chondrogenic gene expression across single cells in distinct cartilage sites.

**(b)** Gene expression matrix and clustering across across cartilage subpopulations identified by SmartSeq2 scRNAseq.

**(c)** Tabulation of subpopulation distribution across cartilage sites.

**(d)** Selected gene expression across cartilage subpopulations showing genes related to cartilage, stem and progenitor cells as well as proliferation.

**(e)** GO Biological Processes analyses based on significantly regulated genes of each cluster compared to all other clusters showing selected significantly enriched processes.

**(f)** RNAScope-based *Panx3* staining at distinct femoral sites showing no expression.

**(g)** Three-dimensional reconstruction of cartilage microexplants from three distinct sites with selected gene expression based on scRNAseq data.

**(h)** Established mouse skeletal stem cell (SSC) lineage tree with cell-surface markers used for prospective identification and isolation.

**(i)** Microarray-based *Panx3* gene expression data of freshly prospectively isolated SSC lineage cell populations from different aged mice (sourced from: GSE34723).

**Extended Data Figure 2: Panx3-expressing cells across mouse and human cartilage tissues. (a)** Re-analyzed mouse femora/tibial epiphyseal tissue scRNAseq data (sourced from:GSE295331). Left: UMAP analysis. Right: Dot plot showing SSC marker and chondrogenic gene expression across unbiased Leiden clusters.

**(b)** *Procr* and *Panx3* gene expression of the same dataset.

**(c)** Re-analyzed human embryonic limb tissue scRNAseq data (sourced from: E-MTAB-14385). Left: UMAP analysis colored by site of origin. Right: Dot plot showing human SSC marker gene expression across skeletal origin sites.

**(d)** Abundance of Panx3^+^ cells across skeletal sites

**(e)** Re-analyzed human adolescent GP scRNAseq data (sourced from: GSE288028). Left: Dot plot showing selected marker gene expression across subclustered GP regions. Right: UMAP based plotting of single cells across GP zones (top) and expression of *PANX3* (bottom).

**(f)** RNAScope-based *Panx3* staining at distinct femoral sites showing no expression.

**(g)** Schematic of Panx3-Cre^ERT2^ mouse generation.

**(h)** Representative image of Panx3-Cre^ERT2^/Ai9 mouse in absence of tamoxifen demonstrating no tdTomato signal leakage.

**(i)** Representative flow cytometric plots of Panx3-Cre^ERT2^/Ai9 mice with and without tamoxifen induction.

**(j)** Quantification of SSC lineage population abundance in tamoxifen-induced Panx3-Cre^ERT2^/Ai9 mice. Data are presented as mean ± SEM; *n* = 6 biologically independent samples.

**Extended Data Figure 3, related to Figure 3: Lineage tracing of Panx3-CreERT2/Ai9-reporter mice in the skeleton. (a)** Representativie section of developing femur bone of Panx3-Cre^ERT2^/Ai9 mice at indicated timepoint.

**(b)** IHC co-staining of tdTomato signal with lineage-specific markers at E10.5 with indicated timepoints.

**(c)** Tamoxifen induction at two distinct timepoints in Panx3-Cre^ERT2^/Ai9 mice showing signal across distinct femoral sublocalizations.

**(d)** Representative femur sections of Panx3-Cre^ERT2^/Ai9 induced at different indicated timepoints and traced for specific periods.

**(e)** Quantification of Panx3-derived tdTomato+ cells after 24 chase at 5-months. *n* = 3 biologically independent samples.

**(f)** IHC co-staining of tdTomato signal with lineage-specific markers at indicated age and timepoints.

**Extended Data Figure 4: Panx3-Cre^ERT2^/Ai9-reporter mice across skeletal sites and in response to skeletal challenges. (a)** Representative image of Panx3-Cre^ERT2^/Ai9 mouse vertebrae induced at 8-weeks of age and analyzed at 14-months.

**(b)** Representative image of Panx3-Cre^ERT2^/Ai9 mouse cranial suture area induced either early in life or upon adulthood as indicated.

**(c)** Representative image of Panx3-Cre^ERT2^/Ai9 mouse mandible tissue induced either early in life or upon adulthood as indicated.

**(d)** Representative image of Panx3-Cre^ERT2^/Ai9 mouse femur with tdTomato co-stained with Col1a1.

**(e)** Representative flow cytometric analysis showing tdTomato+ fraction of microdissected epiphyseal bone are including GP and metaphyseal area with trabecular bone below the GP of Panx3-Cre^ERT2^/Ai9 mouse femurs. Control treatment or PTH1-34 for two weeks are compared.

**(f)** Co-staining of dtTomato and ColX in Panx3-Cre^ERT2^/Ai9 mouse femoral fracture callus tissue.

**Extended Data Figure 5: Panx3-expressing cells are crucial regulators of skeletal formation and repair. (a)** Representative Movat’s Pentachrome stain of developing femurs in control and Panx3-Cre^ERT2^/DTA mice.

**(b)** GP area of control and DTA mice tamoxifen-induced at P1 for six days.

**(c)** Flow cytometric analysis of SSC lineage subpopulations in femurs of DTA mice as indicated. Data are presented as mean ± SEM; *n* = 3-4 biologically independent samples.

**(d)** Representative TRAP staining of metaphyseal region of femurs in control and DTA mice.

**(e)** Representative flow cytometric gating strategy to identify immune cell populations in mouse blood.

**(f)** Representative flow cytometric gating strategy to identify hematopoietic stem cell (HSC) lineage populations in mouse bone marrow. pHSC: phenotypic HSC.

**(g)** Quantification of blood immune cell populations. Data are presented as mean ± SEM; *n* = 4 biologically independent samples. Statistical significance was assessed using a two-tailed unpaired Student’s *t* test. Exact *P* values are indicated in the figure.

**(h)** Quantification of bone marrow HSC lineage cell populations. Data are presented as mean ± SEM; *n* = 4-8 biologically independent samples. Statistical significance was assessed using a two-tailed unpaired Student’s *t* test. Exact *P* values are indicated in the figure.

**(i)** Representative IHC images of control and DTA SSC-derived ossicles in renal capsule. Top: IHC for ColX. Bottom: RNAScope for *Panx3*. Right: IHC for CD11b and corresponding quantification. Data are presented as mean ± SEM; *n* = 3 biologically independent samples. Statistical significance was assessed using a two-tailed unpaired Student’s *t* test. Exact *P* values are indicated in the figure.

**(j)** Fracture experiment schematic using Panx3-Cre^ERT2^/DTA mice (top). Middle left: Representative micro-CT image of 2-weeks fracture callus in control and DTA groups. Middle right and bottom: Micro-CT based analysis of indicated bone parameters of callus. Data are presented as mean ± SEM; *n* = 6-7 biologically independent samples. Statistical significance was assessed using a two-tailed unpaired Student’s *t* test. Exact *P* values are indicated in the figure.

**(k)** Fracture experiment schematic using Panx3-Cre^ERT2^/DTA mice (top). IHC for CD45 and cKit in fracture callus tissue of control and DTA mice (left). Volumetric bone parameters of same mice shown on right. Data are presented as mean ± SEM; *n* = 3-44 biologically independent samples. Statistical significance was assessed using a two-tailed unpaired Student’s *t* test. Exact *P* values are indicated in the figure.

**(l)** Representative IHC images of Col1a1 and Osteopontin (Opn) in microfracture (MF) regenerate in control and DTA mice.

**(m)** Representative IHC co-stainign of Acan, Col1a1, ColX and Opn with dtTomato after MF regeneration.

**Extended Data Figure 6: Introduction of Panx3-expressing SSCs in articular cartilage. (a)** Gene expression of Wnt and Fgf-related signaling factors across cartilage subpopulations identified using scRNAseq analysis.

**(b)** LIANA-based significant ligand-receptor signaling interactions between Panx3^+^ subcluster cells and other clusters.

**(c)** Single cell spatial reconstruction of cartilage microexplants showing *Sdc4* gene expression in individual cells across cartilage sites (top) and ligand-receptor expression of *Bmp-Bmpr* and *Fgf-Fgfr* genes (bottom).

**(d)** Quantification of in vitro chondrogenesis and osteogenesis assays with distinct addition of factors (top) and represented staining images below. Data are presented as mean ± SEM; *n* = 3 biologically independent samples. Statistical significance was assessed by one-way ANOVA followed by Tukey’s multiple-comparisons test. Exact *P* values for the indicated comparisons are shown in the figure.

**(e)** Flow cytometric quantification of BCSP abundance in MF site. Data are presented as mean ± SEM; *n* = 3 biologically independent samples. Statistical significance was assessed by one-way ANOVA followed by Tukey’s multiple-comparisons test. Exact *P* values for the indicated comparisons are shown in the figure.

**(f)** Clonal analysis of MF site in Panx3-Cre^ERT2^/Rainbow mice showing quantification of cells per clone. Data are presented as mean ± SEM; *n* = 3 biologically independent samples. Statistical significance was assessed by one-way ANOVA followed by Tukey’s multiple-comparisons test. Exact *P* values for the indicated comparisons are shown in the figure.

**(g)** Co-stain of tdTomato and ColX or Opn in MF site of factor treated mice.

**(h)** MF experiment with different factor combination in wild type mice analyzed 4-or 8-weeks after procedure. Sectopm stained with Safranin O.

**(i)** Same MF experimenet in 1-year-old wild type mice analyzed 8- and 16-weeks after MF.

**(j)** MF experiment in mice with experimentally induced OA (DMM). Safranin O stained representative section 8-weeks after MF are shown.

**(k)** Representative Safranin O stained section of adult mouse joint with established OA and subsequent MF 20-weeks after MF and treated with Fgf2 and Wif1 at time of injury.

